# Empirical validation of the nearly neutral theory at divergence and population genomic scale using 144 placental mammals genomes

**DOI:** 10.1101/2025.04.17.649326

**Authors:** M. Bastian, D. Enard, N. Lartillot

## Abstract

By limiting the efficacy of selection, random drift is expected to play a major role in genome evolution. Formalizing this idea, the nearly-neutral theory predicts that the ratio of non-synonymous over synonymous polymorphism (*π*_*N*_ /*π*_*S*_) within populations, and divergence (*d*_*N*_ */d*_*S*_) between species, should both correlate negatively with *N*_e_. This has previously been tested in mammals and other groups. However, most studies have focused on either *d*_*N*_ */d*_*S*_ or on *π*_*N*_ /*π*_*S*_, thus not addressing the problem across evolutionary scales. In addition, many studies at the macro scale have used life-history traits (LHT) as a proxy of *N*_e_, assuming that large-bodied organisms have lower *N*_e_ than small-bodied species. However, this assumption itself has rarely been validated against more objective measures of *N*_e_, such as genetic diversity *π*_*S*_ = 4*N*_e_*µ*, in part because *π*_*S*_ estimates are scarce. Here we propose an integrative test of the nearly-neutral predictions on 150 mammalian species, using 6000 orthologous genes, spanning the macro and the micro-evolutionary scale, using for the latter a measure of heterozygosity on each of the assembled diploid genomes. At the micro scale, we observe, for the first time in mammalian nuclear genomes, a relationship between *π*_*N*_ /*π*_*S*_ and *π*_*S*_. At the macro scale, we confirm the positive correlation between *d*_*N*_ */d*_*S*_ and LHT but, more importantly, establish that LHT and *d*_*N*_ */d*_*S*_ are correlated with *π*_*S*_, although weakly so. Together, these results provide the first global test of the nearly-neutral theory in mammals across time scales, suggesting all variables are correlated with a single hidden variable: *N*_*e*_.

**Significance Statement:** Natural selection eliminates large-effect deleterious mutations but has limited efficacy for purging mutations of small effects. The effective population size (called *N*_e_) is what determines the cutoff below which selection is inefficient. Since *N*_e_ varies between species, the prediction is that more mutations of intermediate effect should segregate and reach fixation in species with small populations. This has been widely studied both between species (macro-evolution) and within species (micro-evolution), although never in a fully integrated manner. Here, we analysed 144 genomes of placental mammals and 6002 orthologous genes, using the original sequence reads produced for genome assembly to estimate within-species genetic diversity based on the heterozygosity of the diploid individual whose genome has been sequenced. We show consistent relationships between genetic diversity within species, life-history traits such as body mass or longevity and selection efficacy simultaneously at the micro- and macro-evolutionary levels. Our analysis provides a globally consistent picture confirming the role of effective population size in tuning selection efficacy in placental mammals.

## 1 Introduction

Phylogenomics and population genomics have been the subject of extensive research over the years. However, they are rarely used in tandem or even compared. While macro-evolution is widely viewed as the long-term outcome of micro-evolutionary processes, direct integration of empirical divergence and polymorphism data over a broad set of species is still quite uncommon. Yet, a more decisive integration along those lines would benefit from the complementary strengths of polymorphism and divergence in investigating important aspects of evolutionary processes and their long-term impact on genomes.

One particularly important point on which this comparison could bring important insights is the question of the role of effective population size (*N*_e_), and its variation, in genome evolution. Random drift limits the efficacy of selection, implying that many features of genomes might in fact be neutral or even maladaptive (Lynch and Gabriel, 1990; Lynch and Conery, 2003; Nei *et al*., 2010). The tradeoff between selection and drift was first formalized by Ohta and Kimura (Ohta and Kimura, 1971; Ohta, 1973a), leading to the nearly-neutral theory. According to this theory, the majority of mutations are deleterious, with a broad distribution for their fitness effects (DFE), ranging from slightly to strongly deleterious. In this context, the inverse of the effective population size acts as a threshold on the DFE, such that the fate of mutations with selective effects smaller than 1*/N*_e_ is typically determined by genetic drift and may thus segregate and reach fixation, while the mutations with larger effects are efficiently purified by natural selection (Ohta, 1973a).

In the case of protein coding genes, synonymous mutations are often considered as neutral, at least in groups such as mammals (Ohta, 1995), thus providing a useful neutral background against which to evaluate the fate of non-synonymous mutations, which on the other hand have typically a broad range of fitness effects (Eyre-Walker *et al*., 2006; Eyre-Walker and Keightley, 2007). Contrasting the non-synonymous and synonymous compartments thus yields two measurable quantities: at the population (or micro-evolutionary) scale, the ratio of non-synonymous over synonymous polymorphism (or *π*_*N*_ /*π*_*S*_) and at the phylogenetic (or macro-evolutionary) scale, the ratio of non-synonymous over synonymous divergence (or *d*_*N*_ */d*_*S*_). These two metrics measure the combined effect of selection and random drift. In both cases, lower values indicate that non-synonymous deleterious mutations are more efficiently purged by natural selection and are thus segregating at lower frequency (in the case of *π*_*N*_ /*π*_*S*_) or less prone to reaching fixation (in the case of *d*_*N*_ */d*_*S*_). According to the nearly neutral theory, both are thus expected to decrease as a function of *N*_e_, with a quantitative response that depends on the exact shape of the DFE (Welch *et al*., 2008) or on the structure of the fitness landscape (Lourenço *et al*., 2011; Goldstein, 2013; Latrille and Lartillot, 2021).

Testing these predictions, however, raises the question of how to estimate *N*_e_ and more fundamentally, how to define it (reviewed in Waples, 2022). In particular, some define it from a demographic point of view, when others invoke *N*_e_ as a more inclusive measure of genetic drift (thus including non-demographic contributions, such as linked selection). Some authors argue for a more complex definition of *N*_e_ and derive the concept at different time scales (Nadachowska-Brzyska *et al*., 2022; Müller *et al*., 2022). In the following, we opt for the inclusive definition of *N*_e_, as a net measure of the intensity of drift, as it is more directly relevant for what determines the processes of population genetics and molecular evolution.

Concerning the estimation of *N*_e_, the cases of macro- and micro-evolution are quite contrasted. At the micro-evolutionary scale, genetic diversity provides direct quantitative access to *N*_e_, through the relation *π*_*S*_ = 4*N*_e_*µ* where *µ* is the mutation rate per site, per generation (Charlesworth, 2009; Wright *et al*., 1939). Of note, *π*_*S*_ depends on both *N*_e_ and *µ*. In principle, *µ* can be estimated, which in turns gives the possibility of directly estimating *N*_e_ based on *π*_*S*_ (Eyre-Walker *et al*., 2002; Brevet and Lartillot, 2021; Lynch *et al*., 2023). In practice, it appears that most of the variation in *π*_*S*_ is contributed by the *N*_e_ component (Lynch *et al*., 2023), making *π*_*S*_ itself a reliable proxy of *N*_e_. Translating the nearly-neutral prediction in this context, *π*_*N*_ /*π*_*S*_ is predicted to correlate negatively with *π*_*S*_, which was indeed observed in mammals, although thus far only for the mitochondrial genome (James *et al*., 2017).

At the scale of divergence between species, molecular evolutionary theory does not provide any simple quantity directly related to *N*_e_. To address this problem, much of previous comparative work has relied on the assumption of a correlation between *N*_e_ and life-history traits (named “LHT” in the following). The main argument is that large and long-lived organisms are generally characterized by a lower abundance than small-bodied organisms characterized by a shorter lifespan (Damuth, 1981; White *et al*., 2007). In turn, *N*_e_ is expected to be directly influenced by true population sizes, at least to some extent. By this argument, random drift is expected to be stronger in large and long-living species (e.g. apes) than in short-lived organisms with small body size (e.g. murid, rodents). In this context, the nearly-neutral prediction is that of a positive correlation between *d*_*N*_ */d*_*S*_ and LHT.

Empirically, a positive correlation between *d*_*N*_ /*d*_*S*_ and LHT has repeatedly been observed (Nabholz *et al*., 2013; Popadin *et al*., 2007; Figuet *et al*., 2016; Romiguier *et al*., 2014) and generally interpreted as a confirmation of the nearly-neutral theory. Actually, the literature on this subject is somewhat contrasted, with both positive and negative results, depending on the exact traits and the clade under scrutiny. More fundamentally, in these analyses, the negative relation between LHT and *N*_e_ is invoked for explaining the observed positive correlation between LHT and *d*_*N*_ /*d*_*S*_, but it is most often not itself independently validated against empirical data. A more robust test of the nearly-neutral theory would certainly require to empirically validate this independent assumption, by testing the correlation between LHT and an independent measure of *N*_*e*_. Alternatively, one could measure the correlation of *d*_*N*_ /*d*_*S*_ more directly with this independent measure of *N*_e_, rather than with the indirect proxy represented by LHT.

Since no clear quantitative measure of *N*_e_ is available at the macro-evolutionary scale, an independent validation of the correlation between LHT and *N*_e_ could rely on the micro-evolutionary proxy *π*_*S*_. This has been surprisingly rarely investigated, essentially in metazoans (Romiguier *et al*., 2014) and in birds (Figuet *et al*., 2016) – in the latter case, precisely because the positive correlation between LHT and *d*_*N*_ /*d*_*S*_ was not observed, such that a deeper investigation was felt needed. Correlation analyses between *π*_*S*_ and LHT have never been reported in the case of mammals. Similarly, there have been few attempts of a direct confrontation of *d*_*N*_ /*d*_*S*_ with *π*_*S*_ (Eyre-Walker *et al*., 2002; Brevet and Lartillot, 2021), which have also been limited in statistical power by the use of a small number of data points for *π*_*S*_. Altogether, the current picture of the validity of the nearly-neutral theory across time scales is still incomplete.

The globally incomplete and undecisive picture offered by previous work is further affected by the use of disparate methods for estimating the molecular quantities and testing for their correlations. In this respect, one important aspect is to properly account for phylogenetic non-independence (Felsenstein, 1985), and this, in a context where some of the quantities of interest (in particular *d*_*N*_ /*d*_*S*_) are not, strictly speaking, observed in extant species (as is the case for life history traits or *π*_*S*_ and *π*_*N*_ /*π*_*S*_), but are only indirectly inferred along the branches of the phylogeny.

To deal with this issue, in the previous literature, two main avenues have been explored: a sequential approach, and an integrative one. The sequential approach consists in first estimating *d*_*N*_ /*d*_*S*_ on the terminal branches of the phylogeny and tabulating those estimates along with the other relevant variables. Then this table and a phylogeny are analysed using a standard methods based on independent contrasts (PGLS or Phylogenetic Generalized Least Squares, Pagel, 1997). In brief, these methods assume that the traits of interest jointly evolve according to a multivariate Brownian process over the tree and estimate the variance-covariance matrix of the process by maximum likelihood, based on the observed values at the tips. The integrative approach, on the other hand, proposes to directly integrate the comparative test (the phylogenetic regression) into the phylogenetic codon model used for inferring the patterns of synonymous and non-synonymous substitutions over the tree (Lartillot and Poujol, 2011). In practice, this amounts to condition the codon model on branch-specific rates of synonymous and non-synonymous substitutions that are derived from the underlying multivariate Brownian process, which is otherwise identical to that assumed by the PGLS approach. These two methods have their strengths and weaknesses. However, they have rarely been confronted on the same data (but see Figuet *et al*., 2017).

Finally, in addition to relying on a robust methodology as clarified above, a proper testing of the nearly neutral predictions bridging the two times scales also requires adequate data at the genomic scale and globally across species of a large clade. In this direction, the currently available data from population genomics projects are still relatively scarce, making it difficult to gather data on intra-specific polymorphism for all species across a clade, that would be matched with the genes included in the multiple sequence alignment used for reconstructing the patterns of *d*_*N*_ /*d*_*S*_ over the phylogeny.

On the other hand, the increasing availability of whole genomes sequences in some taxa (Zoonomia, 2020; Feng *et al*., 2020) suggests an alternative approach not depending on population genomics projects. At least for taxonomic groups such as mammals, one can rely on the fact that all published genomes are from diploid individuals. As such, they contain information about the heterozygosity of those individuals, which itself is informative about the genetic diversity of the population (Li and Durbin, 2011). Although published genome sequences typically do not provide this information, it is nevertheless possible to obtain it by returning to the original sequencing reads (Figuet *et al*., 2016; Zoonomia, 2020). Compared to a true population genomics project, this way to proceed provides more limited information, potentially affected by the fact that some individuals may happen to be inbred. On the other hand, it gives access to intra-specific diversity for all species for which complete genomes are available. Thus, it makes it possible to obtain a nearly complete data matrix, genome wide and over an entire clade, with which intra and inter-specific information can be globally confronted in the context of a comparative analysis.

In this article, we propose to combine whole-genome orthologous genes and heterozygosity data across a wide range of mammals. For this purpose, we annotated the complete one-to-one orthologous genes from 144 whole genomes. We then did variant calling on them to obtain the heterozygosity on the annotated genes for all species that are present in the phylogeny. This heterozygosity can then be confronted with the rich information currently available for mammals about life history traits (De Magalhaes *et al*., 2009). We performed our correlation study using both an integrative and sequential approach, to do a global test of the nearly-neutral theory.

## 2 Results

Starting from 197 complete mammalian genomes and after several steps of data processing and filtering, we gathered a dataset of 6002 single copy orthologous genes for 144 placental species. Species and genes were chosen to ensure sufficient assembly quality, and maximise completeness of the data matrix. The genes were aligned and trimmed, and a phylogeny was reconstructed using a sample of 1000 of these genes. Based on this reference dataset, we considered an additional species subsampling scheme, meant to reduce the noise contributed by non-phylogenetic variation between closely related species. Specifically, we trimmed the dataset to keep only species that are separated by at least 10 My of divergence, resulting in a dataset of 89 species (referred to as the sparse dataset in the following, as opposed to the original dense dataset of 144 species).

This provides the macro-evolutionary backbone of our analysis, which was complemented with information about intra-specific polymorphism. Specifically, for each species represented in the dataset, the original reads that were used for genome assembly were mapped onto the genome to detect heterozygous sites. We focussed on the SNPs that were called in the coding sequences of the 6002 genes. This yields a set of synonymous and non-synonymous SNPs from which a *π*_*S*_ and *π*_*N*_ /*π*_*S*_ were inferred.

Out of the 144 species of the dense phylogenetic dataset, 6 had an abnormal allele frequency distribution and were removed for all subsequent analyses, thus giving 138 species with data about synonymous and non-synonymous heterozygosity. Visual inspection of the relation between *π*_*S*_ and *π*_*N*_ /*π*_*S*_ (Figure S1) suggests that 6 additional species, all of which have low coverage, may have insufficient data for reliable heterozygosity estimation. For this reason, we decided to implement a refined version of the dataset in which the polymorphism of these six species was excluded from the analysis. Here we analyze this adjusted version of the data (thus with 132 species with data about synonymous and non-synonymous heterozygosity). Results that include the six outlier species are presented in the supplementary material (Figure S12). In the context of the sparse phylogenetic backbone, this translates into 82 (or 86 including the 6 outliers just mentioned) species with data about heterozygosity, out of the 89 species represented in the phylogeny. All species filtering and subsampling steps are summarized in Supplementary Material, Table 5.11.

The relation between *π*_*S*_, *π*_*N*_ /*π*_*S*_, *d*_*N*_ /*d*_*S*_ and LHT were then examined using both an integrative method, FastCoevol, adapted from the Coevol approach Lartillot and Poujol (2011) and a sequential approach based on PGLS.

### 2.1 Micro-evolutionary scale: *π*_*S*_ **versus** *π*_*N*_ /*π*_*S*_

At the micro-evolutionary scale, the efficacy of selection is reflected by *π*_*N*_ /*π*_*S*_, while *π*_*S*_ provides a direct proxy of *N*_e_. The nearly-neutral theory predicts a negative correlation between these two measures. Plotting *π*_*N*_ /*π*_*S*_ against *π*_*S*_ (Figure S1), a clear negative correlation is indeed observed between these two variables, except for six species (the hazel dormouse *Muscardinus avellanarius*, the hirola *Beatragus hunteri*, the cotton rat *Sigmodon hispidus*, the black rhinoceros *Diceros bicornis*, the humpback dolphin *Sousa chinensis* and the mantler howler *Alouatta palliata*) showing a abnormally low *π*_*S*_, given their *π*_*N*_ /*π*_*S*_. This could be the sign of recent inbreeding. However, these six species also tend to show a low genomic coverage and, for some of them, a poor genome assembly (see Material and Methods), which suggests more a data quality issue for these outlying points rather than a low real *π*_*S*_.

Excluding these six low *π*_*S*_ species, a clear log-linear negative relation between *π*_*S*_ and *π*_*N*_ /*π*_*S*_ is observed (Figure 1), which is confirmed by the phylogenetic correlation analysis (Figure 2). The correlation is also significant and present similar correlation coefficient when these six species are included (Figure S12). This is compatible with a more efficient purging of slightly deleterious mutations (lower *π*_*N*_ /*π*_*S*_) in species with large *N*_e_ (larger *π*_*S*_), in agreement with the nearly-neutral prediction.

**Figure 1.**
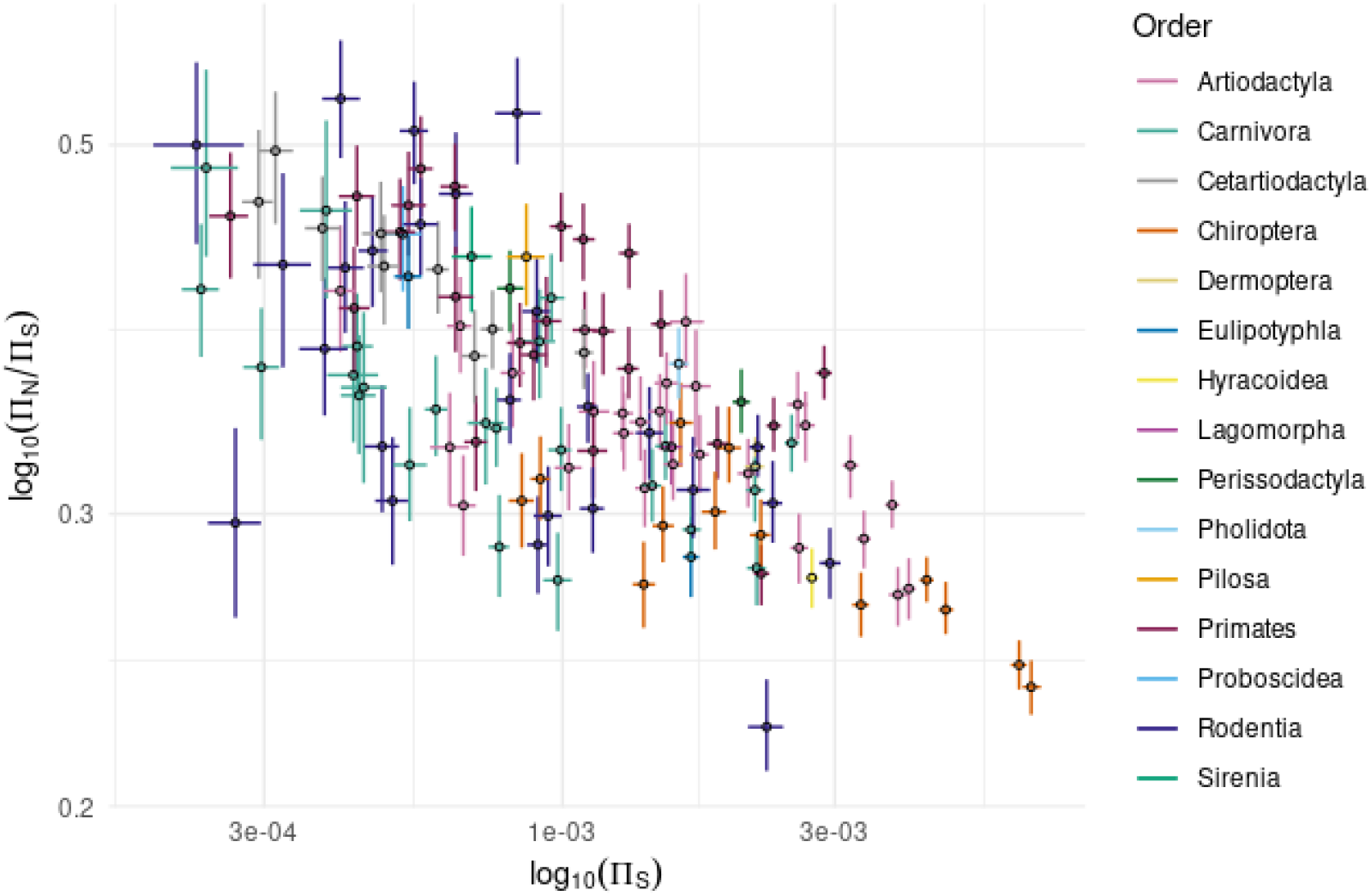
**Plot of** *π*_*N*_ /*π*_*S*_**against** *π*_*S*_**(in log-log scale)** for all species of the analysis except the six species with less than 1000 SNPs (bars: 95% CI computed by non-parametric bootstrap). Colours correspond to species families.

**Figure 2.**
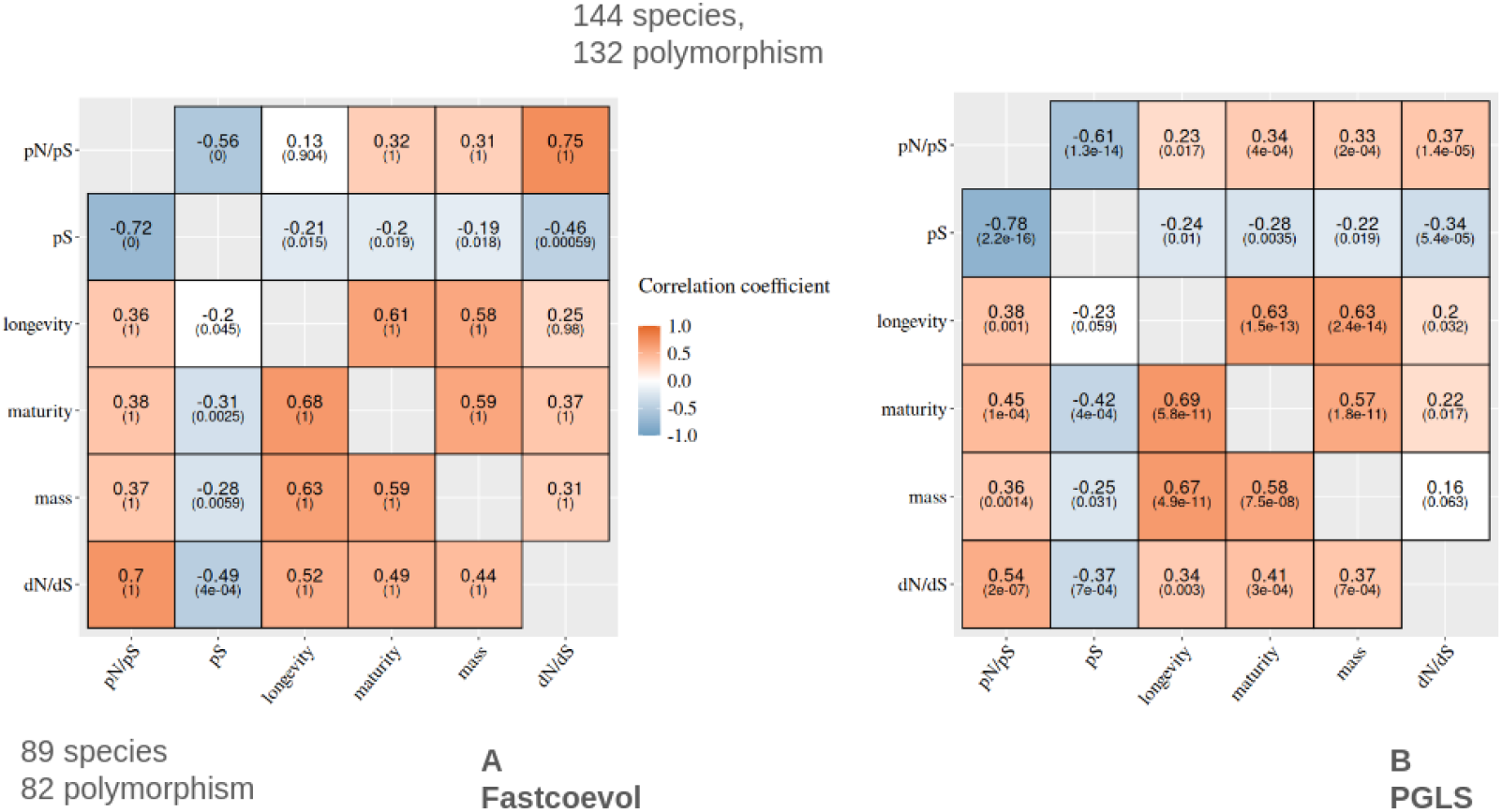
Correlation coefficients from the FastCoevol and PGLS analyses. **A**: FastCoevol analysis. **B**: PGLS analysis. **Upper diagonal**: using the 144-species phylogeny. **Bottom diagonal**: using the 89-species phylogeny. Colours correspond to the magnitude of the correlation coefficients (*r*). Numbers in brackets correspond to the posterior probability for FastCoevol analyses (which must be higher than 0.975 or lower than 0.025 for the correlation to be considered significant) or p-value for PGLS analyses (which must be lower than 0.05 for the correlation to be considered as significant). Non-coloured estimates correspond to non-significant correlations.

Of note, we observe a stronger correlation between *π*_*S*_ and *π*_*N*_ /*π*_*S*_ with the sparse (89 species) phylogenetic dataset than with the original dense version with 144 species (*r* = *−*0.72 versus *r* = *−*0.56 with FastCoevol, *r* = *−*0.78 versus *r* = *−*0.61 with PGLS, Figure 2, compare above and below the diagonal in each panel). It may seem paradoxical that removing data points increases the strength of the inferred correlations. A reasonable explanation is that the independent contrasts between closely related species are dominated by estimations errors or short-term effects and that reducing the number of species reduces this source of noise.

The slope of log *π*_*N*_ /*π*_*S*_ as a function of log *π*_*S*_ is estimated at -0.16, with 95% credibility interval (-0.20, -0.13). This is smaller in absolute value than what was found in mammalian mitochondrial genomes (of the order of -0.63, James *et al*. (2017)), or in primates (-0.29, Brevet and Lartillot (2021)).

### 2.2 Macro-evolutionary scale: life history traits versus *d*_*N*_ /*d*_*S*_

At the macro-evolutionary scale, life history traits are known to be strongly correlated with each other across mammals. This is recovered in our analysis using both PGLS and FastCoevol (Figure 2). As for *d*_*N*_ /*d*_*S*_, its history over the phylogeny such as inferred by FastCoevol is shown in Figure 3 which presents the *d*_*N*_ /*d*_*S*_ reconstruction along the 89-species tree. From this figure, we observe a *d*_*N*_ /*d*_*S*_ ranging from 0.13 to 0.28 over the tips. Taxonomic groups with the lowest *d*_*N*_ /*d*_*S*_ are Rodentia (i.e. *Peromyscus maniculatus*, the eastern deer mouse, *d*_*N*_ /*d*_*S*_ = 0.139) and Lagomorpha (i.e. *Ochotona princeps*, the american pika, *d*_*N*_ /*d*_*S*_ = 0.13). At the other end of the range, taxonomic groups with the highest *d*_*N*_ /*d*_*S*_ correspond to Cetacea (i.e. *Eschritius robustus*, the gray whale, *d*_*N*_ /*d*_*S*_ = 0.285) and Primates (i.e. *Alouata palliata*, the mantled howler *d*_*N*_ /*d*_*S*_ = 0.222). This general observations from Figure 3 is globally suggestive of a pattern of a high *d*_*N*_ /*d*_*S*_ in large and long lived mammals (also recovered using the 144 species tree S5).

**Figure 3.**
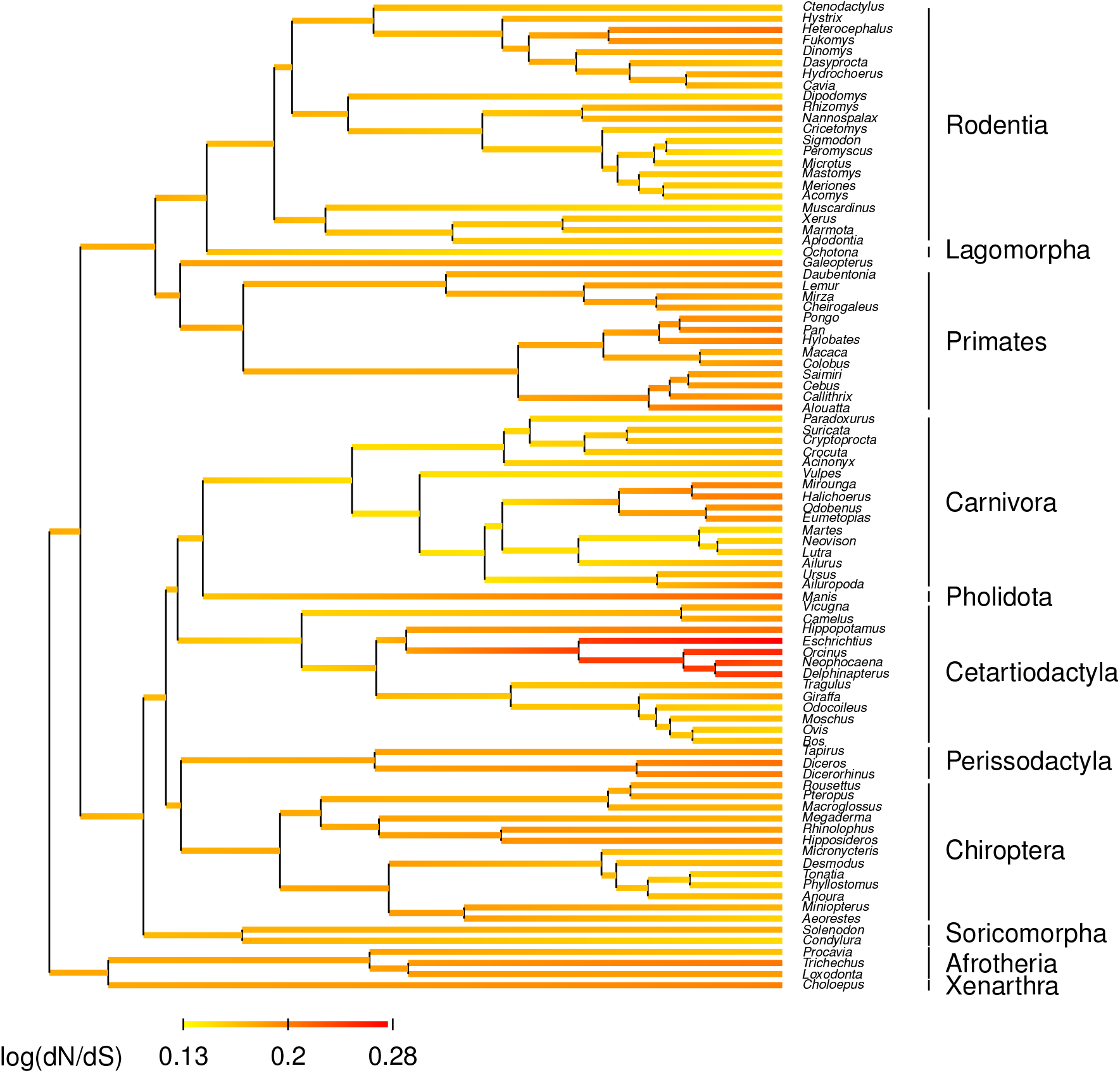
Reconstruction of log(*d*_*N*_ /*d*_*S*_) along the phylogeny of placental mammals (89 species) by FastCoevol.

On the dense (144 species) tree, the FastCoevol analysis shows a significant positive correlation between *d*_*N*_ /*d*_*S*_ and all life history traits, with correlation coefficients varying from *r* = 0.25 to 0.37 depending on the trait (Figure 2 A, upper diagonal). Similarly, the PGLS analysis shows positive correlations (*r ≃* 0.21), all significant except for the correlation between *d*_*N*_ /*d*_*S*_ and body mass (Figure 2 B, upper diagonal). The correlations are slightly stronger with FastCoevol, which can be explained by the fact that the additional variance contributed by the short-term overdispersion of *d*_*N*_ /*d*_*S*_, which is modelled by FastCoevol, has been discounted. Using the sparse (89 species) tree, we observe stronger correlations under both methods. With the sparse dataset, correlations between *d*_*N*_ /*d*_*S*_ and mass also become significant in the PGLS analysis. This effect is also presents in the FastCoevol analysis, which suggests that the short-term over-dispersion of *d*_*N*_ /*d*_*S*_ along branches is not totally captured by the overdispersion component of the integrative approach.

Altogether, both PGLS and FastCoevol analyses agree on a significant positive correlation between *d*_*N*_ /*d*_*S*_ and LHT at the macro-evolutionary scale. These results are generally interpreted as an indirect correlation of both variable with *N*_e_ and are thus compatible with a nearly-neutral interpretation (Ohta, 1973b, 1992) suggesting less efficient selection in long-lived, large-bodied mammals, presumably characterised by lower *N*_e_ over the tree. As it stands, however, this interpretation is reliant on the additional and thus far untested assumption that LHT indeed correlate with *N*_e_, a point which we now investigate.

### 2.3 Confronting the micro- and macro-evolutionary scales

If macro-evolutionary patterns are the long term result of micro-evolutionary processes, we expect to observe correlations of our metrics of interest across timescales. However, this depends on the extent to which shortterm *N*_e_ (such as measured by *π*_*S*_) reflects long-term *N*_e_, and to what extent long-term *N*_e_ is indeed reflected in life history traits.

In this direction, with FastCoevol, we found significant negative correlations between *π*_*S*_ and life history traits, although only in the analysis without the six low-*π*_*S*_ species. Even then, the correlation coefficients are weak and below the significant threshold for longevity in the 89 species analysis (Figure 2A, bottom diagonal). The PGLS analysis also shows weak correlations and the statistical support vary depending on the life history traits and the taxon sampling (Figure 2B).

On the other hand, *π*_*N*_ /*π*_*S*_ shows a robust negative correlation with life history traits, for the sparse and dense datasets and with the two methods. The most straightforward explanation for this correlation is that both variables are indirectly correlated with *N*_e_. The weaker correlation between *π*_*S*_ and LHT, compared to that between *π*_*N*_ /*π*_*S*_ and LHT, could be due to normalization errors in *π*_*S*_ and *π*_*N*_ estimates, which would cancel in their ratio (see discussion).

Here also, as above for *d*_*N*_ /*d*_*S*_ versus life history traits, the correlations are stronger and more consistent using FastCoevol rather than PGLS and when the dataset is thinned to remove closely-related species. Again, this could be due to the account for short-term overdispersion of *d*_*N*_ /*d*_*S*_ performed by FastCoevol and to the negative impact of non-phylogenetic variation. In the present case, short-term fluctuation in *N*_e_ may contribute. Taken together, these correlation patterns of *π*_*S*_ and *π*_*N*_ /*π*_*S*_ with life history traits, which are congruent in their sign, but contrasted in their strength and significance, are consistent with the hypothesis of a connection between shortand long-term Ne.

Conversely, and finally, *d*_*N*_ /*d*_*S*_ correlates negatively with *π*_*S*_, thus providing a more direct evidence of an effect of *N*_e_ on the efficacy of selection at the macro-evolutionary scale. The slope of log *d*_*N*_ /*d*_*S*_ as a function of log *π*_*S*_ is equal to *−*0.07(*−*0.12, *−*0.04). It is thus significantly shallower than that of log *π*_*N*_ /*π*_*S*_ as a function of log *π*_*S*_ (with non-overlapping credible intervals). The *d*_*N*_ /*d*_*S*_ also correlates positively with *π*_*N*_ /*π*_*S*_. Overall, the correlation coefficient are globally stable across the different analyses for *d*_*N*_ /*d*_*S*_ versus *π*_*S*_, but more variable for *d*_*N*_ /*d*_*S*_ versus *π*_*N*_ /*π*_*S*_.

## 3 Discussion

The nearly neutral theory states that selection is less efficient at purging mildly deleterious mutations in smaller populations. Accordingly, it predicts higher levels of segregating non-synonymous polymorphisms within species and higher rates of non-synonymous substitutions between species, relative to the synonymous compartment, when *N*_e_ is low (Ohta, 1973b). In this study, we aim to confirm this hypothesis at two time scales: micro-evolutionary (using *π*_*S*_ and *π*_*N*_ /*π*_*S*_) and macro-evolutionary (using life history traits and *d*_*N*_ /*d*_*S*_), based on a large dataset of 6002 genes in 144 placental mammals.

The correlations observed in our study are recapitulated in Figure 4. Overall, all correlations are globally compatible with a nearly neutral interpretation. More specifically, all could be explained by all variables being correlated with a single hidden variable, which is *N*_e_. The results are robust to alternative subsampling schemes, except for the correlation between *π*_*S*_ and life history traits, which is perhaps the weakest and least convincing among all observed correlations.

**Figure 4.**
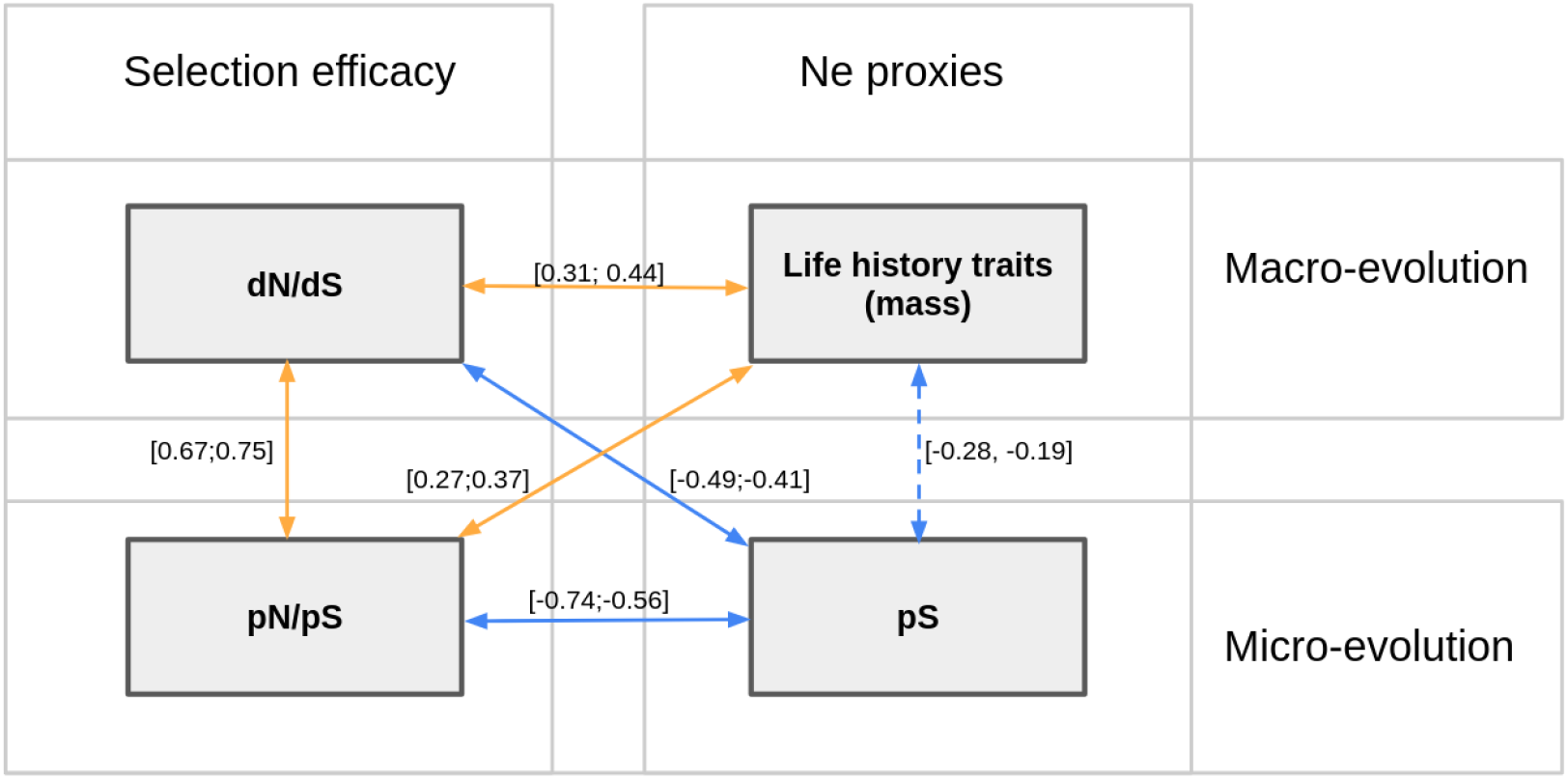
Schematic summary of the correlations observed in the present study using FastCoevol. Negative correlations are in blue, positive correlations in orange. Values within brackets correspond to the range of correlation coefficients obtained under the four different analyses using FastCoevol (sparse and dense dataset, with and without low *π*_*S*_ species). Mass is used here to represent life history traits. The dotted line between life history traits and *π*_*S*_ represents the instability of the relationship, which depends on the dataset used.

**Figure 5.**
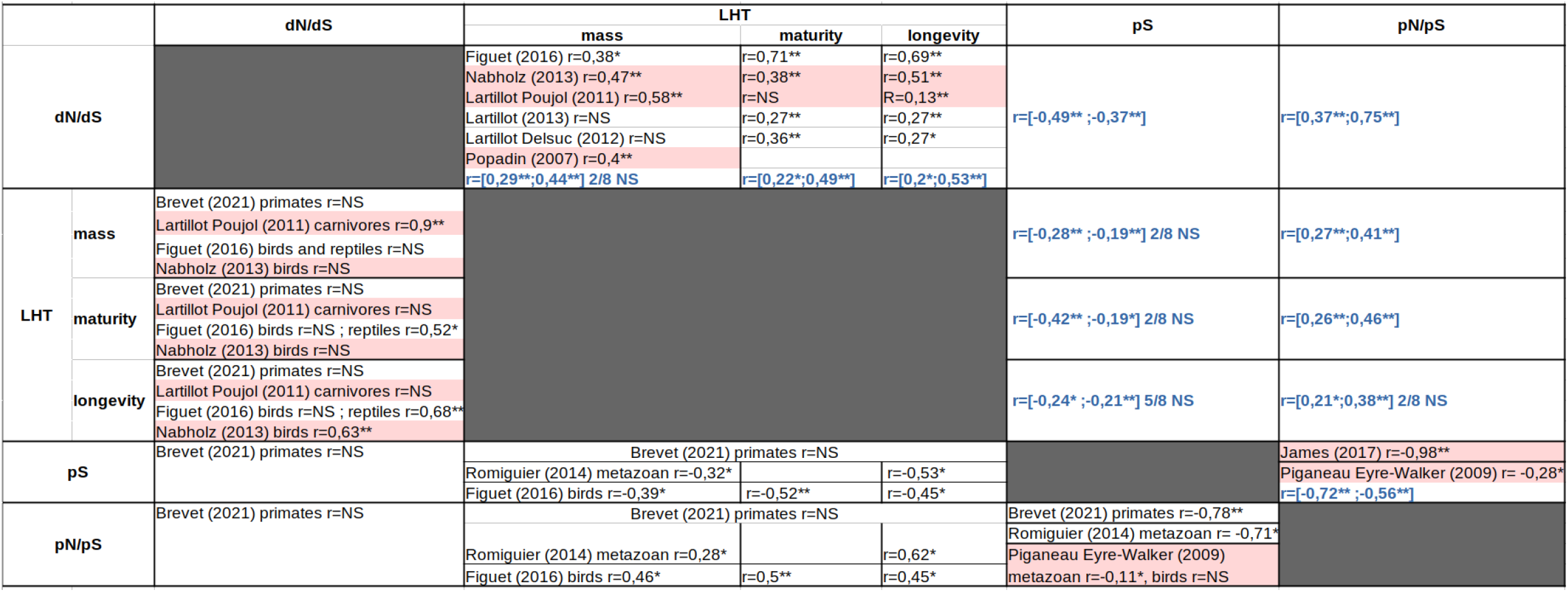
Table summarizing previously published results. Upper diagonal: Studies on mammals. Lower diagonal: Studies on other taxonomic groups. “*” mean a significant correlation (p-value *<* 0.05, pp*>* 0.95 or pp*<* 0.05), “**” strongly significant correlation (p-value *<* 0.001, pp*>* 0.975 or pp*<* 0.025); “NS”: non-significant correlation. Results in bold blue correspond to our study. The values are within brackets as they represent the min and max of the correlation coefficient from the eight alternative sampling and method scheme. When a correlation is not significant in one of the eight modalities, we emphasize it by giving the number of non statistically significant observations over the eight. Results in pink correspond to analysis using mitochondrial data.

As presented in the introduction, several prior studies have already examined some of these correlations. In Table 5, we present a summary from the published literature concerning both the significant correlations and those sought but not found to be significant (labelled as “NS”). These different projects are not easy to compare as they are not consistent in data and methods used. Our results are also reported in this table, which thus highlights the correlations that we found and that were missing until now. Among them, the most important ones are: first, the correlation between LHT and *π*_*S*_, which, even if it is weak, gives more objective support to the belief that LHT reflect *N*_e_ in mammals; second, the correlation between *d*_*N*_ /*d*_*S*_ and *π*_*S*_, which provides a more direct test of a nearly-neutral effect on long-term protein evolution than the now well-established correlation between *d*_*N*_ /*d*_*S*_ and LHT; and finally, the correlation between *π*_*N*_ /*π*_*S*_ and *π*_*S*_, which provides an equivalent test of the nearly-neutral predictions at the micro-evoluationary scale, and which was previously established only in the case of mitochondrial genes in mammals (James *et al*., 2017) and was still missing for the nuclear compartment.

### 3.1 Limitations and potential sources of error

All the successive steps in our pipeline, from genome annotation to the alignment of gene sequences and the reconstruction of a phylogeny, are subject to different types of errors which could accumulate and could impact our conclusions (Simion *et al*., 2020). Moreover, genomes of poor quality could be particularly problematic. Given the variable quality of the genome assemblies used here, a lot of work has been done to control for data errors (See Material and Methods).

Nevertheless, some errors may still be present in some amount. It thus raises the question of whether they could drive or compromise some of the correlations observed here. Different sources of errors at micro and macro-evolutionary scale, which are susceptible to bias the correlation analyses, can be identified.

#### Errors in alignment and SNP calling

At macro-evolutionary scale, alignment errors tends to inflate the apparent *d*_*N*_ /*d*_*S*_. This effect is stronger for short branches (measured in synonymous evolutionary divergence) which could create an artefactual correlation between *d*_*N*_ /*d*_*S*_ and *d*_*S*_. If long-living species tend to have a low *d*_*S*_, this could induce an artefactual correlation between *d*_*N*_ /*d*_*S*_ and life history traits. Similarly, at micro-evolutionary scale, SNP calling errors will tend to inflate the apparent *π*_*N*_ /*π*_*S*_. This effect will be stronger in species with a low true *π*_*S*_ and can create an artefactual correlation between *π*_*S*_ and *π*_*N*_ /*π*_*S*_. By extension, if long-living mammals tend to have a low *π*_*S*_, this will induce an artefactual correlation between *π*_*N*_ /*π*_*S*_ and life history traits. It is still difficult, as it stands, to completely rule out the possibility that some of the correlation that we observe here are driven by these potential biases. Of note, concerning *π*_*N*_ /*π*_*S*_, this would imply a strong correlation between *π*_*S*_ and life-history traits.

#### Estimation of the number of mutational targets and its impact on *π*_*S*_and *π*_*N*_

To compute the synonymous and non-synonymous rates at the two scales, we need to normalise the counts by the number of mutational targets. In the case of polymorphism, this in turn depends on the estimation of the number of callable positions. Here, the number of callable positions is not readjusted after the filtering of low quality SNPs. This is particularly an issue concerning lower quality genomes because they present less raw SNPs detected by variant calling and lower SNP quality overall, inducing a stronger reduction of the total number of final SNPs, without removing them from the callable set. Of note, this bias should affect *π*_*S*_ and *π*_*N*_ but should cancel in their ratio, and thus is not expected to impact *π*_*N*_ /*π*_*S*_. Such normalisation problems could explain the fact that *π*_*S*_ has much weaker correlation than *π*_*N*_ /*π*_*S*_ with the other traits.

#### Segregating polymorphism and its impact on *d*_*N*_ /*d*_*S*_

Segregating polymorphism implies that the apparent *d*_*N*_ /*d*_*S*_ on the terminal branches of the tree contains a certain proportion of *π*_*N*_ /*π*_*S*_ in its estimation. This could contribute to making the apparent correlation between *π*_*N*_ /*π*_*S*_ and *d*_*N*_ /*d*_*S*_ stronger than it actually is. Indeed, this correlation is among the strongest observed in our analysis. More generally, this suggests care in interpreting the correlation of *d*_*N*_ /*d*_*S*_ with other variables, as it cannot be excluded that they entail a hidden correlation with *π*_*N*_ /*π*_*S*_. On the other hand, this effect cannot explain everything – in particular, it cannot explain the fact that the correlation between *d*_*N*_ /*d*_*S*_ and LHT is stronger than that between *π*_*N*_ /*π*_*S*_ and LHT.

#### Use of heterozygosity of an individual genome as a proxy for population diversity

The use of the heterozygosity of a single individual as a proxy for population-level diversity has several conceptual and practical limitations. First, even if the heterozygosity computed on a whole genome is supposed to represent the population polymorphism, it can still deviate substantially from the population mean for a lot of reasons (inbreeding, population structure, bottleneck *etc*). Second, individual heterozygosity is influenced by genome-wide stochasticity. Even in a large, panmictic population, the realized heterozygosity of one diploid genome can differ from the expectation simply by chance, and this variance remains unquantified. Third, technical factors such as sequencing depth, variant calling errors, and assembly fragmentation can disproportionately affect heterozygosity estimates when only one genome per species is available. These limitations imply that heterozygosity from a single individual may not exactly capture the polymorphism of the population. However, in our study, both *π*_*S*_ and *π*_*N*_ /*π*_*S*_ are derived from the same individual, so their correlation is robust to these sources of variance even if their absolute values may not precisely reflect the real *π*_*S*_ and *π*_*N*_ /*π*_*S*_.

Altogether, our results are still subject to several limitations and remain to be confirmed, in particular by implementing a more robust normalisation of *π*_*S*_ and by quantifying the contribution of the polymorphism to the *d*_*N*_ /*d*_*S*_ estimates. Nevertheless, as it stands, at worse the relation between *π*_*S*_ and LHT is underestimated, and the relation between *π*_*N*_ /*π*_*S*_ and LHT is perhaps the most robust result reported here pertaining to the relationship between micro and macro evolution.

### 3.2 Quantitative scaling of *π*_*N*_ /*π*_*S*_ and *d*_*N*_ /*d*_*S*_ as a function of *π*_*S*_ and *N*_e_

The slope of the relationship between *π*_*N*_ /*π*_*S*_ and *π*_*S*_, or that between *d*_*N*_ /*d*_*S*_ and *π*_*S*_ (on a log-log scale), have been of some interest in the previous literature, owing to their bearing on the shape of the underlying distribution of fitness effects. Specifically, if the DFE is gamma, of shape parameter *β*, and in the limit where *β* is small, then the slope of log *π*_*N*_ /*π*_*S*_ or *d*_*N*_ /*d*_*S*_ as a function of log Ne is expected to be equal to *−β* (Welch *et al*., 2008). Of note, this assumes that the DFE is constant across species, a point which has been questioned, both empirically (Castellano *et al*., 2019) and theoretically (Goldstein, 2013; Latrille *et al*., 2021). Also, as discussed in Brevet and Lartillot (2021), the variation of *µ* between species, which tends to be opposite to that of *N*_e_, can result in the slope of log *π*_*N*_ /*π*_*S*_ as a function to log *π*_*S*_ being smaller that that of log *π*_*N*_ /*π*_*S*_ as a function of log *N*_e_.

Here we see that the slopes as a function of *π*_*S*_ are globally shallower than previously reported results for mitochondria across mammals for *π*_*N*_ /*π*_*S*_ (James *et al*., 2017) or nuclear genomes in primates for both *π*_*N*_ /*π*_*S*_ and *d*_*N*_ /*d*_*S*_ (Brevet and Lartillot, 2021). This could be the result of different behaviors in different genomic compartments or of clade-specific effects. Alternatively, incorrect estimation of *π*_*S*_ (see above) could also contribute to making apparent slopes shallower than true slopes.

We also observe a shallower slope for *d*_*N*_ /*d*_*S*_ than for *π*_*N*_ /*π*_*S*_ (as in Brevet and Lartillot (2021), but now with non-overlapping credible intervals). As discussed in Brevet and Lartillot (2021), this could be due to a non-nearly neutral contribution to the *d*_*N*_ /*d*_*S*_ (e.g. positive selection), unrelated to *N*_e_. Alternatively, a random short-term effect on *N*_e_, on the top of its long-term phylogenetic trends, is expected to push the slope of *d*_*N*_ /*d*_*S*_ (but not *π*_*N*_ /*π*_*S*_) towards smaller values.

### 3.3 Perspectives on micro versus macro-evolution

Our analysis provides a decisive entry point for further exploration of the micro-macro connections. About the methods, in the end, the use of an integrative (FastCoevol) rather than a sequential (PGLS) method doesn’t seem to make a big difference. They both gives similar results. However, integrative methods can be furthered developed to reconstruct *N*_e_. In particular, in this article, we make the assumption that *π*_*S*_ represents *N*_e_, using the relation *π*_*S*_ = 4*N*_e_*µ*. However, *µ* itself varies between species, and is known to correlate with life history traits (Lanfear *et al*., 2014; Lynch *et al*., 2023). Correcting for *µ* would therefore be important in order to have a direct access to *N*_e_. A classical approach to correct for *µ* is to estimate it by using the synonymous divergence between close species and fossil calibration with a generation time (Eyre-Walker *et al*., 2002). In the integrative method, we can incorporate a decomposition of *π*_*S*_ into *N*_e_ and *µ* components by the reconstruction of *µ* along the topology. This was already implemented in Brevet and Lartillot (2021) and could be recruited and applied to the dataset obtained here for mammals.

Using this approach would make it possible to estimate slopes directly with respect to *N*_e_, not *π*_*S*_. However, this would first require explicitly modelling the mismatch between short-term and long-term *N*_e_. Such an improvement in the model would provide a more quantitative idea about the intensity of short term fluctuations of *N*_e_ and see in particular whether it explains the shallower slope for *d*_*N*_ /*d*_*S*_, compared to *π*_*N*_ /*π*_*S*_.

Finally, here we focused on nearly neutral evolution, but our dataset could be used for investigating other aspects, as it presents the opportunity to compare two genomic compartments which respond differently to certain evolutionary processes such as positive selection (McDonald and Kreitman, 1991) or mutational bias and genetic biased conversion toward GC (Duret and Galtier, 2009).

## 4 Materials and Methods

Figure 6 summarizes the pipeline developed and used here, from the acquisition of the data to the final analyses. Below, we describe in detail the different steps of this pipeline.

**Figure 6.**
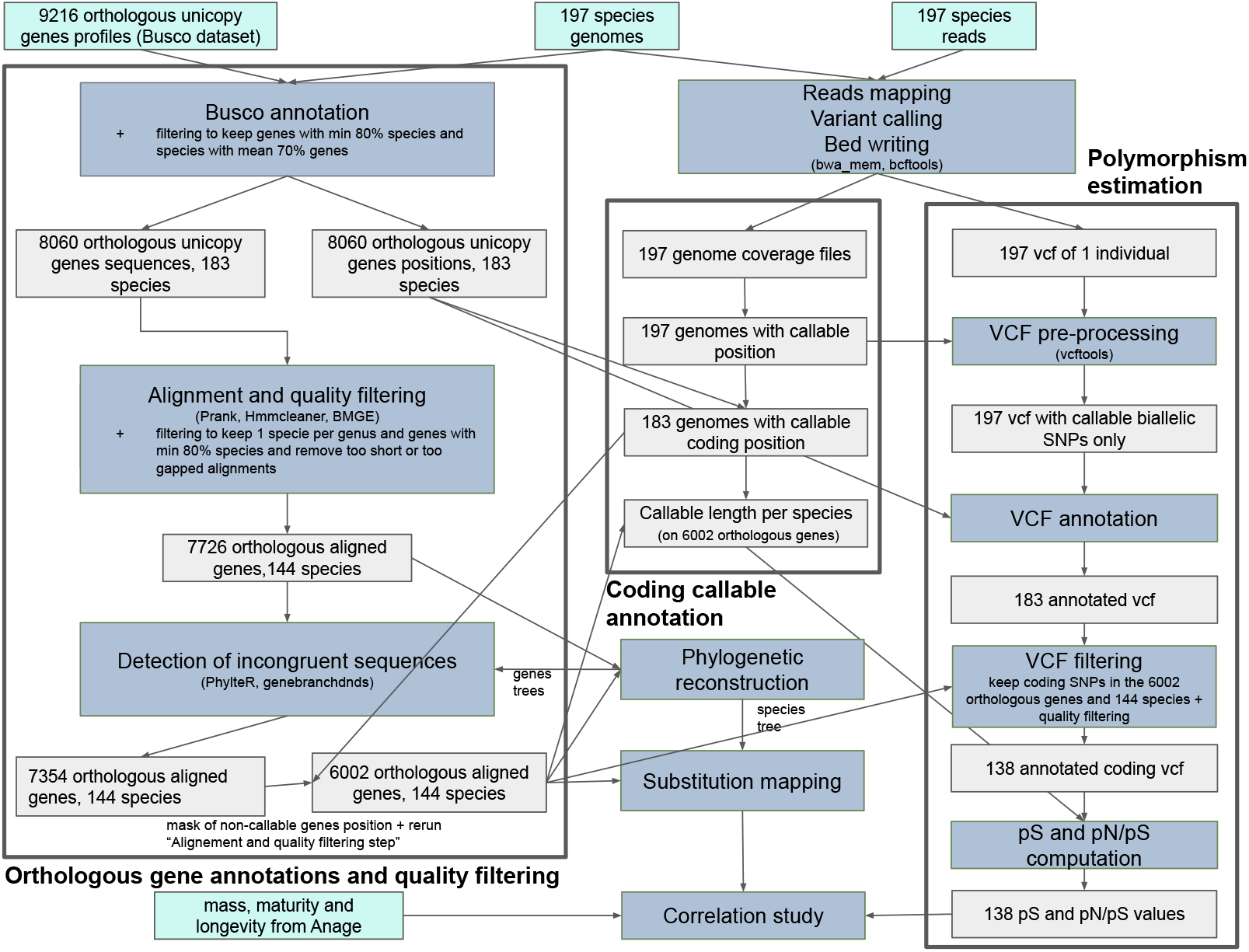
Simplified summary of the pipeline from whole genomes acquisition and vcf writing to the final correlation analysis. Light blue boxes correspond to the input data, grey boxes to the intermediate data and blue boxes to different steps of the pipeline. A more complete and more detailed desciption of the pipeline is available at https://github.com/MeloBastian/Mam_Ne

### 4.1 Data acquisition

Four types of input data were used (depicted in light blue in Figure 6): complete genomes in fasta format, their associated reads, a BUSCO database and life-history traits. The reference genome assemblies were downloaded from NCBI Genomes in July 2021. At this time, we selected the assemblies of eutherian mammals using different criteria. First, we selected assemblies with median contig size (N50) of at least 30kb to avoid overly truncated genes. We then select assemblies with at least one individual with sequencing depth higher than 20x according to NCBI Genomes statistics. In addition, we excluded the assemblies from domestic species whose genetic variation is well-known to be reduced by the domestication process and the lab individuals that are usually highly inbred.

For each of the 197 assemblies that met all these requirements, we selected the latest reference assembly available in July 2021. Because the entire analysis relies on heterozygosity in single individuals, when multiple individuals were sequenced for a genome assembly project, we only selected and downloaded the reads from the individual with the highest sequencing depth. Sequencing reads were then mapped on their corresponding assembly with bwa mem2 v2.2.1 used with default parameters (Vasimuddin *et al*., 2019). We then conducted a variant calling to identify the heterozygotes positions and create for each species a whole-genome sequencing depth files, which was very useful for the sequences filtering.

### 4.2 Orthologous gene annotation and filtering

We developed a pipeline starting from the 197 assembled complete genomes, producing in output a data matrix of aligned and filtered single copy orthologous genes. At different steps of the process, quality controls and filters were applied (detailed below), leading to the removal of some species or genes from the analysis. Additionally, closely related species can share ancestral polymorphism. This shared polymorphism can affect the estimation of the *d*_*N*_ /*d*_*S*_. To mitigate this issue, we chose to retain only one species per genus. The reduced dataset at the end of the pipeline contain 144 species and 6002 single-genes alignments (genes number per species and other metrics are resumed in Supplementary material, table 5.11).

#### 4.2.1 Identification of orthologs

Not all genomes are annotated, and those that are were annotated using different procedures. Since our focus is on orthologous coding sequences, and to ensure a homogeneous treatment across the entire species set, we annotated all genomes using BUSCO (Simão *et al*., 2015). BUSCO is a commonly used bioinformatics tool for assessing genome assembly quality based on the identification of orthologous genes assumed universal across the taxonomic group specified. Here we relied on the mammalian database (orthodb.10), defining a set of 9226 orthologous unicopy genes across this clade. The program provides a percentage of recovered orthologous genes, which is meant as an indicator of the genome quality. BUSCO also provides intermediate output files specifying the coordinates of the orthologous genes identified in the genome being analysed. These intermediate files were processed so as to generate, for each genome, a fasta file containing all coding sequences and a bed file with their position along the genome. From our initial dataset of 197 species, we only kept species with BUSCO complete single-copy score higher than 70%, meaning that at least 70% of the searched genes were found. In a second step, we removed genes present in less than 80% of the species to ensure their universality across placental mammals. After this step, we are left with 183 mammalian species and 8060 BUSCO one-to-one orthologous complete genes.

#### 4.2.2 Alignment and quality filtering

The sequences of each orthologous gene were aligned using PRANK (Löytynoja, 2014). To remove poorly aligned regions that could bias the phylogenetic inference, we filtered them using HMMcleaner (Di Franco *et al*., 2019) and BMGE (Criscuolo and Gribaldo, 2010) with default parameters. These two methods are complementary and improve the global quality of the alignments. BMGE trims unreliably aligned columns based on gap patterns, while HMMCleaner identifies and removes short, species-specific segments that deviate from the profile HMM of the alignment (Ranwez and Chantret, 2020).

Upon filtering, some alignments ended up empty or with a lot of gaps. We removed orthologous genes for which the alignment has less than 10 nucleotides or more than 75% gaps, resulting in 8056 aligned fasta files. Next, our dataset was reduced to only one species per genus leading to a set of 144 species. When multiple species occur for a genus, the one with the higher whole-genome coverage was chosen. We then removed the genes with less than 80% occurrence in this species set, resulting in a list of 7726 genes.

#### 4.2.3 Detection of incongruent gene using gene tree

Although BUSCO aims at identifying single-copy orthologs, some alignments may nevertheless contain sequences with incorrect orthology assignments. Gene-level alignments for which the true evolutionary history differs from that of the species may introduce noise (Boussau and Scornavacca, 2020) and have a strong adverse input on the study and in particular on the species-level *d*_*N*_ /*d*_*S*_ estimates. Other sources of errors may also be present, such as sequencing errors resulting in compensated frameshifts, incorrect exon identification, etc.

To identify such deviant sequences and genes, we used a combination of two approaches. First, we reconstructed the 7726 gene trees using Iqtree2 with defaults parameters (Minh *et al*., 2020). The trees where then given as an input to PhylteR (Comte *et al*., 2023), which analyses all gene trees and provides a list of sequences and genes exhibiting outlier behaviour. Second, we developed a Bayesian shrinkage model (see Supplementary Material section 5.12) to identify sequences showing large deviations from the global pattern of synonymous branch lengths across the tree, which are expected to be very similar across genes.

PhylteR detected 20 genes that required complete removal and 1565 genes that contain at least one sequence to be removed. The Bayesian shrinkage model identified 1695 genes with at least one sequence displaying a deviant synonymous branch length. In total, 2015 genes out of the initial 7726 were flagged for at least one sequence, by either PhylteR or the Bayesian shrinkage model, with 1074 genes identified by both methods. We removed all the flagged sequences from the alignments and also genes containing less than 80% of the species after this filtering step, resulting in a dataset set of 7354 gene alignments.

#### 4.2.4 Callable fraction of the genome

Using genomes and their associated reads mapped onto them, we computed bam files containing the sequencing depth at each position of the genomes using bcftools v1.13 (Danecek *et al*., 2021). Given the large size of the resulting files, we converted the sequencing depth per position into a sequencing depth per window of 100 bp. We then computed for each species a mean genome sequencing depth and identified the 100 bp windows for which sequencing depth is two times higher or lower than this mean. These sites were considered as non-callable and were masked for the rest of the analysis. One species, *Neotragus schauinslandi*, for which more than 80% of the genome was masked, was removed from the analysis. Without *Neotragus schauinslandi*, the mean non-callable fraction of the genome is around 12% with a maximum of 40%.

Of note, some of the genomic regions considered as non-callable according to the criteria just described may overlap the coding sequence contained in the multiple sequence alignment across placentals (see above). To address this potential issue, we came back to the 7354 genes obtained after the use of PhylteR and the Bayesian shrinkage model, and replaced the non-callable coding sites by gaps directly in their Prank output. We then reran the filtering pipeline described above from HMMCleaner to the exclusion of the too short or gapped sequences and of the genes represented in less than 80% of species. This resulted in a final high-quality dataset of 6002 genes with few missing data.

### 4.3 Alternative species subsampling schemes

To investigate the impact of closely related species in our correlation analysis, we subsampled the 144 species to 89 species as follows. Using the dated tree reconstructed by the integrative analysis on the 144 species, we identified all the subclades whose most recent common ancestor is younger than 10% of the total tree depth (corresponding to approximatively 10 My, assuming a date of 100 My for the most recent common ancestor of placental mammals) and chose only one species per subclade, prioritizing species with higher sequencing depth and with all three life history traits informed. This leads to a reduced dataset of 89 species (listed in Supplementary Material). The 6002 single-genes alignments where restricted to these 89 species without realigning. At this step, we cannot remove genes present in less than 80% of species without removing too many genes.

### 4.4 Phylogenetic reconstruction

For the 89 and 144 species dataset, the species tree was estimated using Iqtree2 (Minh *et al*., 2020) with the GTR+G4 model, on a concatenation of 1000 genes. The trees are in Supplementary Material (Figure S6 and S7). For gene trees, required for Phylter (Comte *et al*., 2023), we use Iqtree2 too with default parameters and model search options.

### 4.5 VCF annotation and polymorphism

Variant calling was performed using default parameters of bcftools v1.13 (Danecek *et al*., 2021) (https://samtools.github.io/bcftools/howtos/variant-calling.html). We obtained VCF files containing all variant sites detected. We used vcftools (Danecek *et al*., 2011) to keep only bi-allelic SNPs and remove the SNPs located in the non-callable regions such as previously defined.

SNPs were annotated as synonymous, non-synonymous or non-coding, using a custom Python script that maps the SNPs and the coding orthologous annotation on the genomes. The script also labels each SNP by the name of the gene it belongs to. We then removed the non-coding SNPs and the ones not in the restricted list of 6002 orthologous genes. Furthermore, SNPs with GQ <150 and QUAL < 125 were removed as well as SNPs with an allelic frequency higher than 0.8 or lower than 0.2. After this step, we decided not to further consider the VCF of six species because they present a global abnormal allelic frequency distribution: *Acomys cahirinus* (Cairo spiny mouse), *Przewalskium albirostris* (Thorold’s deer), *Mastomys coucha* (Southern multimammate mouse), *Litocranius walleri* (Gerenuk),*Cheirogaleus medius* (Fat-tailed dwarf lemur) and *Cephalophus harveyi* (Harvey’s duiker) (see Supplementary Material, Figure S4). A summary of the number of SNPs per species at each step is provided in Supplementary Material table 5.11.

After all these filters, six species ended up with less than 1000 SNPs: *Muscardinus avellanarius* (Hazel dormouse), *Beatragus hunteri* (Hirola), *Sigmodon hispidus* (Hispid cotton rat), *Diceros bicornis* (Black rhinoceros), *Sousa chinensis* (Indo-Pacific humpback dolphin) and *Alouatta palliata* (Mantled howler). These species also have a low genome quality based either on the N50 and L50 metrics or on the genome coverage. Below, we will show correlation analyses with and without these 6 species to evaluate their impact on the analysis. As we observe difference in the results using these two dataset, we systematically report the results with and without these six particular species (see Supplementary Material, section 5.9)

Estimates of *π*_*S*_ and *π*_*N*_ were obtained by counting the total number of synonymous (*S*) and nonsynonymous SNPs (*N*) and then normalizing by a rough estimate of the number of synonymous and non-synonymous mutational opportunities (see Supplementary Material 5.2 for a more refine estimate). Thus, denoting by *L* the total number of coding and callable position across all genes, *K*_*S*_ and *K*_*N*_ the number of heterozygote synonymous and non-synonymous positions, and assuming that point mutations in coding sequences are twice more likely to be non-synonymous than synonymous:

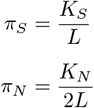

We then compute *π*_*N*_ /*π*_*S*_ using our *π*_*S*_ and *π*_*N*_ measures (reported in Supplementary Material table 5.11).

The sampling variance of our *π*_*S*_ and *π*_*N*_ /*π*_*S*_ estimates was estimated by bootstrap at the gene level (i.e. drawing 6002 genes from the list with replacement and recomputing *π*_*S*_ and *π*_*N*_ /*π*_*S*_). A 95% confidence interval was computed.

We also question the independence between *π*_*S*_ and *π*_*N*_ /*π*_*S*_ witch is itself constructed using a *π*_*S*_ and *π*_*N*_ from a same set of genes (i.e. the independence between A and 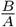). Results are presented in Supplementary Material, Section 5.7.

### 4.6 Life history traits

Data about adult body mass, longevity and age at female sexual maturity (called “maturity” for short in the following) across mammals were obtained from the Anage database (De Magalhaes *et al*., 2009). To mitigate lack of data for some species, we systematically compute the mean of the trait per genus, even if the trait exists for the species (discussed in Supplementary material 5.8). We obtain respectively for the 144 and 89 species datasets, 127 and 81 data points for mass, 118 and 73 for maturity and 118 and 74 for longevity.

### 4.7 Phylogenetic correlation analysis

Phylogenetic correlation analyses between life history traits and molecular variables such as *π*_*N*_ /*π*_*S*_ or *d*_*N*_ /*d*_*S*_ (here after, collectively referred to as “traits”), were conducted using two alternative methods, both of which account for phylogenetic inertia.

#### 4.7.1 Bayesian integrative approach: FastCoevol

Coevol (Lartillot and Poujol, 2011) is an integrative Bayesian approach for analysing the joint evolutionary patterns over the phylogeny of observable traits (such as body mass) and substitution rates (here, *d*_*S*_, *d*_*N*_ /*d*_*S*_). The underlying model recruits the conceptual basis of the independent contrasts approach (Felsenstein, 1985), by modeling the evolution of continuous characters (traits) via a multivariate Brownian diffusion process, while coupling it with a Brownian molecular clock for *d*_*S*_ and *d*_*N*_ /*d*_*S*_ (rates). The coupling implied by the joint

Brownian process for traits and rates enables gene alignments to be incorporated into the study, along with life-history traits and data about heterozygosity in extant species. It provides the opportunity to estimate their joint correlation patterns, while automatically accounting for phylogenetic inertia. Fitting the model to the data produces as an output a dated tree, an estimate of the covariance matrix between traits, as well as a reconstruction of traits along the branches of the tree (see Figure 7).

**Figure 7.**
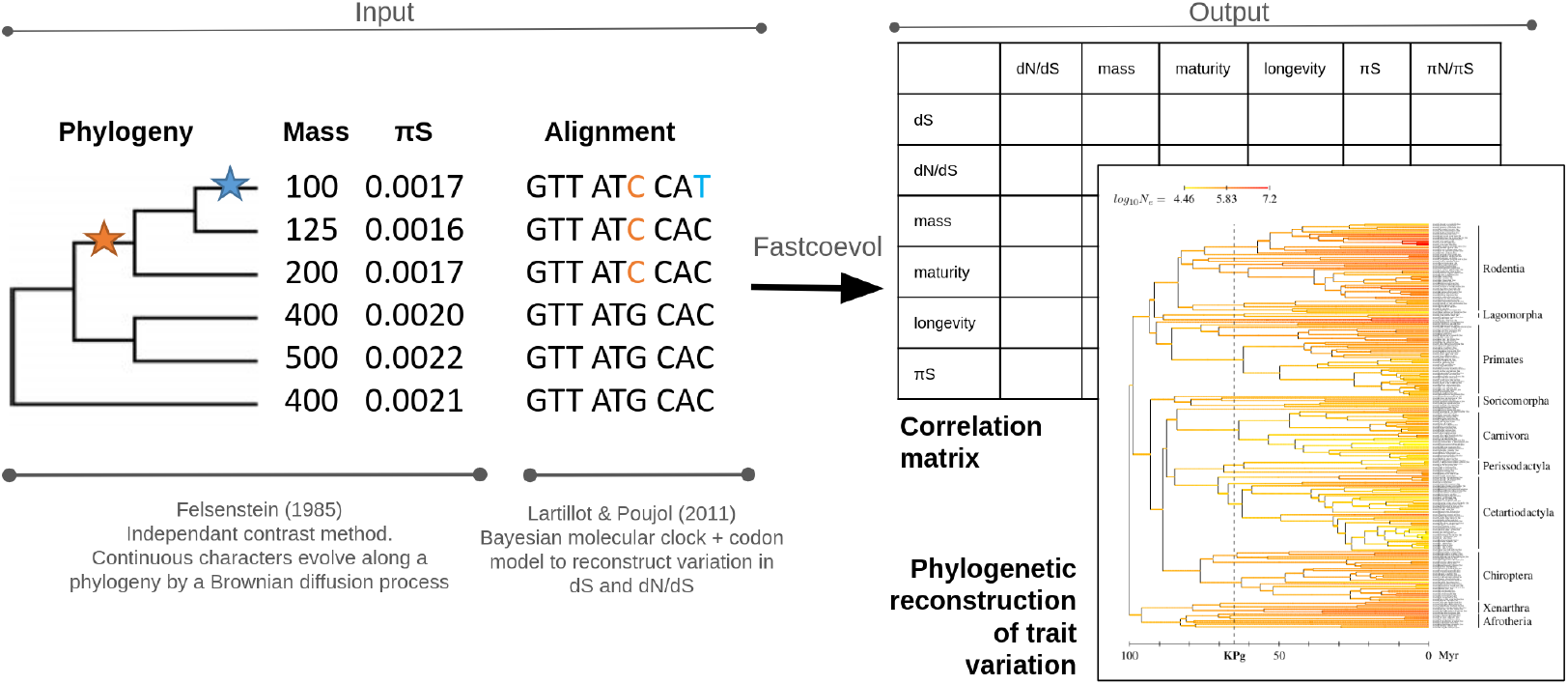
FastCoevol pipeline. The pipeline takes as an input a rooted phylogeny, a table containing the continuous traits of interest (here mass and *π*_*S*_) and a multi-gene alignment. The output is a correlation matrix, with correlation coefficients and posterior probabilities for each pair of traits, including *d*_*S*_ and *d*_*N*_ /*d*_*S*_, and a phylogenetic reconstruction of the variation of each trait along the phylogeny.

For the need of the present project, we implement FastCoevol, a faster version of the Coevol model. This version entails an additional level of branch-specific gamma deviates, meant to account for short-term deviations in *d*_*S*_ and *d*_*N*_ /*d*_*S*_ from the long-term trends implied by the Brownian process. This two-level model is a direct generalization of the mixed relaxed clock model introduced in Lartillot *et al*. (2016) in the context of a codon model.

In its original version, Coevol is computationally intensive, precluding applications on empirical data sets with more than a few tens of genes. To overcome this limitation, FastCoevol is based on substitution mapping approximation (Romiguier *et al*., 2012; Lemey *et al*., 2012). A detailed presentation of the approach is given in the Supplementary Methods (section Bayesian methodology). Specifically, the detailed substitution history for all genes is inferred under a simpler reference model (assuming a constant gene-level *d*_*N*_ /*d*_*S*_ and shared synonymous branch lengths across genes). This substitution history is then collapsed into posterior mean numbers of synonymous and non-synonymous substitutions, and posterior mean numbers of synonymous and non-synonymous mutational opportunities, and this, separately for each branch of the tree. Those numbers are then used as an approximate sufficient statistic for the molecular component of the multivariate Brownian model of Coevol. The use of this computation reduces the computational to 1-2 days for the substitution mapping and less than an hour for the fitting of the Brownian model on the data over the 6002 genes alignments. For computational reasons, we perform the substitution mapping under the reference model on lists of 1000 genes. We then summed statistics across all gene sets, on a per-branch basis, and used this as an input for the FastCoevol model. With FastCoevol, we produced five independent MCMC runs per datasets, for a total of 30000 points. We used a burn-in of 10000 points and computed the posterior mean estimates for the parameters of interest using 1 every 5 points after the burn-in. The values of the five resulting estimates of the covariance matrix were averaged, even though they are very similar to each other. We considered a positive or negative correlation as significant when the posterior probability of the correlation is respectively higher than 0.975 or lower than 0.025, as in Lartillot and Poujol (2011).

#### 4.7.2 Sequential approach: PGLS

As an alternative to the integrative approach of FastCoevol, we used a sequential approach based on the standard PGLS method (Pagel, 1997). Specifically, the mapping statistics just described were summed over all genes and then used to compute an empirical *d*_*N*_ /*d*_*S*_ for each branch. Then, a linear regression analysis with correction for phylogenetic inertia was conducted using a phylogenetic covariance matrix derived from the evolutionary relationships among the species such as inferred by IQTree. For the PGLS analysis, we consider a correlation significant when the p-value is lower than 0.05.

## Acknowledgements

Damien de Vienne (phylter), Laurent Duret (discussions), Julien Joseph (vcf annotation script and discussion), Thibault Latrille (discussion). **Funding:** Agence Nationale de la Recherche (ANR-20-CE02-0008-01 “NeGA”)

David Enard is funded by NIH MIGMS grant 5R35GM142677

## Author contributions

Original idea: N.L M.B; Model conception: N.L; Code: M.B, N.L; Data analyses: M.B, D.E ; Interpretation: M.B, N.L; First draft: M.B; Editing and revisions: M.B, N.L, D.E; Project management and funding: NL.

## Data and materials availability

For scripts:https://github.com/MeloBastian/MamNeFordata:https://doi.org/10.5281/zenodo.15283024

