## supplementary material for "Empirical validation of the nearly neutral theory at divergence and population genomic scale using 144 placental mammals genomes"

### 5 Supplementary Material

#### 5.1 $\pi_S$ and $\pi_N/\pi_S$ bootstrap

This figure is similar to figure 1 in the main article, except that the six species with less than 1000 SNPs are included, so as to graphically display their deviation from the global trend. Point colors stand for sequencing depth, which highlights the low coverage of these six particular species/

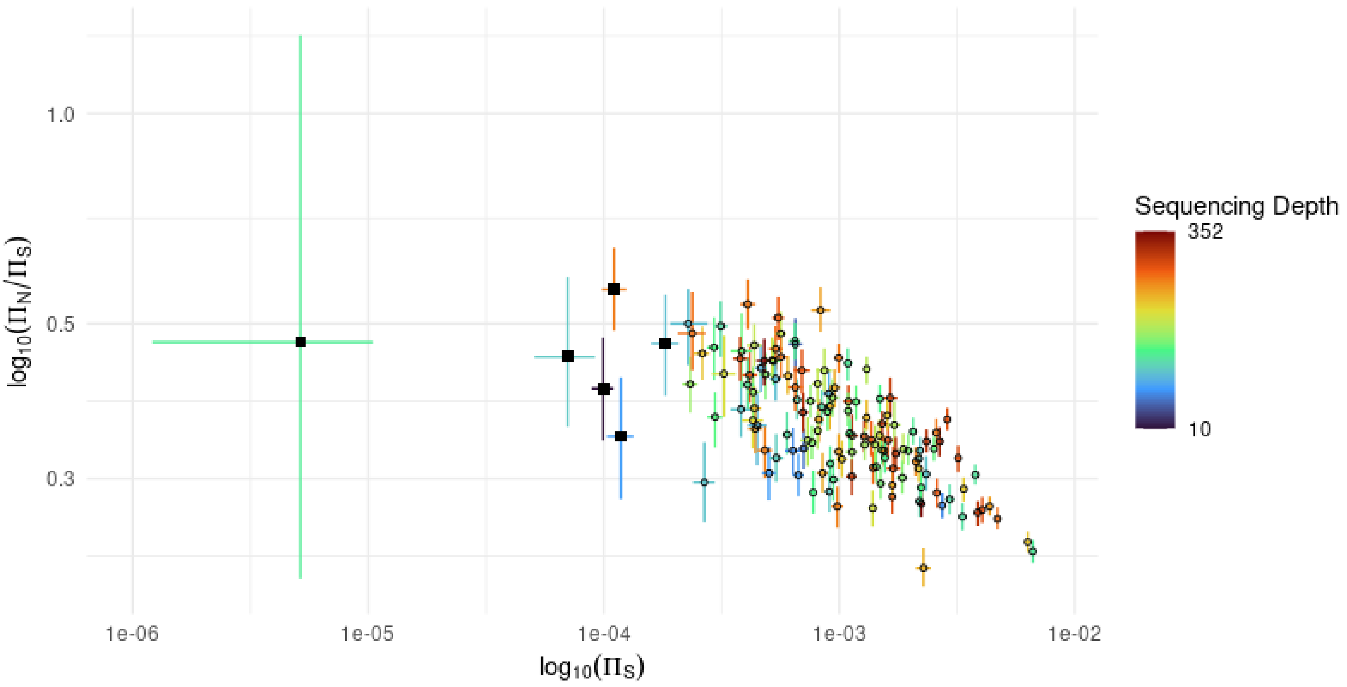


Figure S1: Plot of $\pi_N/\pi_S$ against $\pi_S$ (in log-log scale) for all species of the analysis (bars: 95% CI computed by non-parametric bootstrap). Colours correspond to genome sequencing depth. The square points correspond to the six species with less than 1000 coding SNPs after filtering.

#### 5.2 Estimation of the number of mutational targets for $\pi_S$

In the main article, we estimated the number of synonymous and non-synonymous targets for the $\pi_S$ and $\pi_N$ measures using an approximate method, assuming that the first and second positions in coding sequences are mostly non-synonymous, while the third position is mostly synonymous. This results in an approximate 3:1 ratio.

On the other hand, in our estimation of $d_N$ and $d_S$, we used a codon model to determine with precision the number of synonymous and non-synonymous target in the sequences.

Thus, to align our estimation for $\pi_S$ and $\pi_N$ with that used for $d_N$ and $d_S$, we have implemented a new function in the phylogenetic component of our pipeline, which directly returns the number of mutational targets implied at each tip of the phylogeny by the mutational process assumed by the phylogenetic model starting from the observed sequences. Using this approach, we observe a ratio of 28:77 for synonymous versus non-synonymous targets, instead of the 33:66 ratio used in the simpler method. The difference is substantial, but the ratio is essentially constant across species, and thus this merely shifts the $\pi_S$ and $\pi_N$ estimates by a constant factor, which does not change their correlation patterns (Figure S2, upper diagonal).

As it stands, however, even this approach does not address the possibility that the variation of the transition/transversion ratio between species could affect our estimation of the number of targets, and therefore of $\pi_S$ and $\pi_N$. This is because the phylogenetic model itself, in the version used thus far, assumes a constant mutational process over the tree. To address this, we have implemented a new version of the phylogenetic model which allows for a varying transition/transversion ratio across the tree. We used this new model together with the procedure described above to get yet another more refined estimate of $\pi_S$ and $\pi_N$. With this last method, the correlations inferred for $\pi_N/\pi_S$ with other variables are in fact stronger than with the previous method (Figure S2, lower diagonal). So, in the end, this improved estimation reinforces our conclusions.

| | ds | dN/dS | mass | maturity | longevity | pS | pN/pS |
|---|---|---|---|---|---|---|---|
| dS | | -0,284 (0,01) | -0,58 (0) | -0,734 (0) | -0,653 (0) | 0,129 (0,87) | -0,18 (0,066) |
| dN/dS | -0,394 (0,0006) | | 0,314 (1) | 0,367 (1) | 0,255 (0,98) | -0,455 (0) | 0,751 (1) |
| mass | -0,593 (0) | 0,458 (1) | | 0,594 (1) | 0,584 (1) | -0,192 (0,019) | 0,311 (1) |
| maturity | -0,725 (0) | 0,511 (1) | 0,61 (1) | | 0,607 (1) | -0,194 (0,019) | 0,315 (1) |
| longevity | -0,663 (0) | 0,343 (1) | 0,596 (1) | 0,616 (1) | | -0,208 (0,01) | 0,133 (0,9) |
| pS | 0,119 (0,84) | -0,526 (0,0006) | -0,246 (0,0048) | -0,24 (0,01) | -0,245 (0,0048) | | -0,562 (0) |
| pN/pS | -0,249 (0,016) | 0,812 (1) | 0,414 (1) | 0,207 (1) | 0,207 (0,97) | -0,591 (0) | |

Figure S2: **Correlation coefficient from the FastCoevol analysis with more accurate $\pi_S$ and $\pi_N/\pi_S$ computation.** On the upper side, the transition/transversion ratio in constant over the tree (as in the main results). On the lower side, the transition/transversion ratio is allowed to vary across the tree. Values within brackets correspond to the posterior probabilities, and colours correspond to a significant correlation (pp>0.975 or pp<0.025). The dataset used correspond to the dataset with 144 species, excluding the polymorphism data for the six low $\pi_S$ species.

As for the possible reason why accounting for variation in transition/transversion should lead to stronger correlation patterns: we plotted the transition/transversion ratio measured on external branch against life-history traits and observed a weak positive relation, meaning that the transition/transversion ratio varies across mammals with a higher value for larger mammals (Figure S3). Since a higher transition/transversion leads to a higher proportion of synonymous targets, this would in fact result in a downward bias in the $\pi_N/\pi_S$ estimates for long living species, which would go against the nearly-neutral effect otherwise predicted on $\pi_N/\pi_S$ and would thus partially interfere with it.

##### 5.3 VCF filtering

We annotated the 144 VCF files with a Python script that relates the coding exon positions of the non-filtered 7726 genes from the Busco annotation pipeline to the coordinates in the VCF files and in the genome assembly. We then obtained the coding synonymous, coding non-synonymous or non-coding nature of the SNPs, the name of the corresponding Busco gene and some quality measures for each SNP. We focused on the GQ and QUAL metrics. The QUAL metrics gives the probability that a site has no variant (a false heterozygote). The GQ metric gives the probability that a variant is incorrect (the nucleotide assigned is

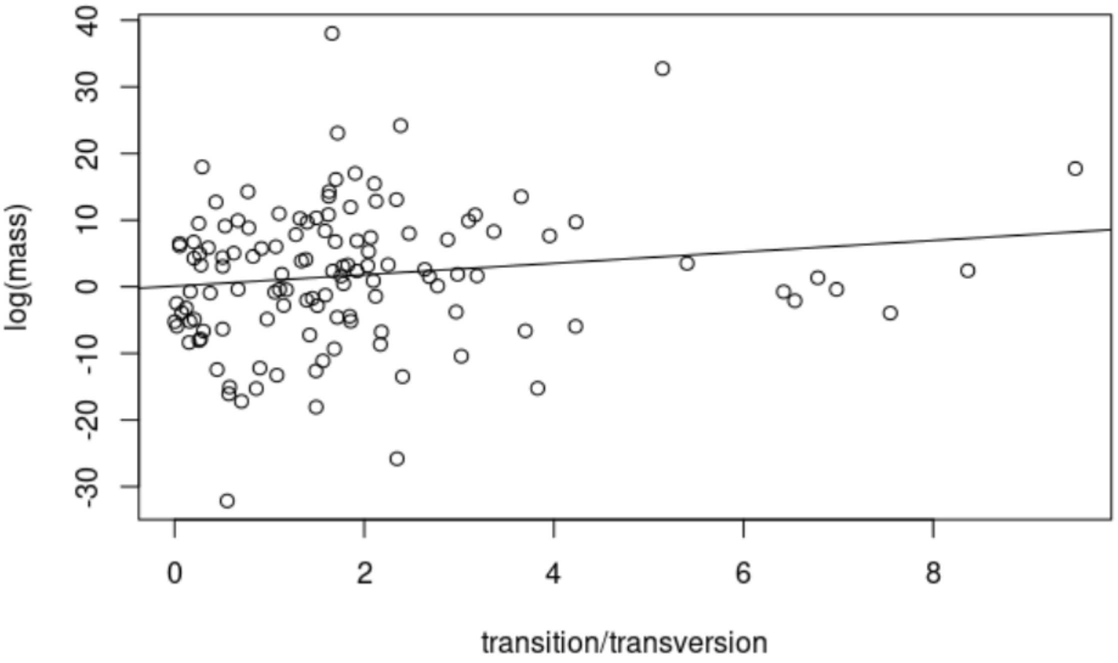


Figure S3: Plot of the external branches transition/transversion ratio computed on a subset of 1000 genes against log(mass) including a phylogenetic inertia correction. Correlation coefficient = 0.21 and p-value = 0.01492

wrong, but there is heterozygosity). These two metrics are useful to our need to have a true variant site with a good call. We applied a strong filtering on the VCF by the use of these metrics (QUAL> 125 and GQ> 150) and applied it to all species, regardless of their genome or calling quality, even if some of them end up with very few SNPs (6 species have less than 1000 SNPs: *Muscardinus avellanarius*, *Beatragus hunteri*, *Sigmodon hispidus*, *Diceros bicornis*, *Sousa chinensis* and *Alouatta palliata*).

The VCF files also provides the number of "reference" and "alternative" reads per SNP. In the case of this study, the reference genome correspond to the individual on which we perform variant calling. The "reference" allele assignment is simply the first read seen in the call and doesn't have any real meaning. We therefore randomized the reference alleles and then computed the frequency of the reference allele. We expect the frequency distribution to be unimodal with a pic around 0.5. We removed the SNP with a frequency lower to 0.2 or higher to 0.8. We graphically inspected this distribution of allelic frequency and detected six species with a distribution (in blue in the Figure S4) different from the expected unimodal one centered around 0.5 (in red in the Figure S4). The six species are *Acomys cahirinus*, *Przewalskium albirostris*, *Mastomys coucha*, *Litocranius walleri*, *Cheirogaleus medius* and *Cephalophus harveyi*. These six VCFs are removed from the analysis. Figure S4 shows an example of a species with an allelic balance distribution that matches with our expectation and two species with an abnormal distribution in two different manners. s

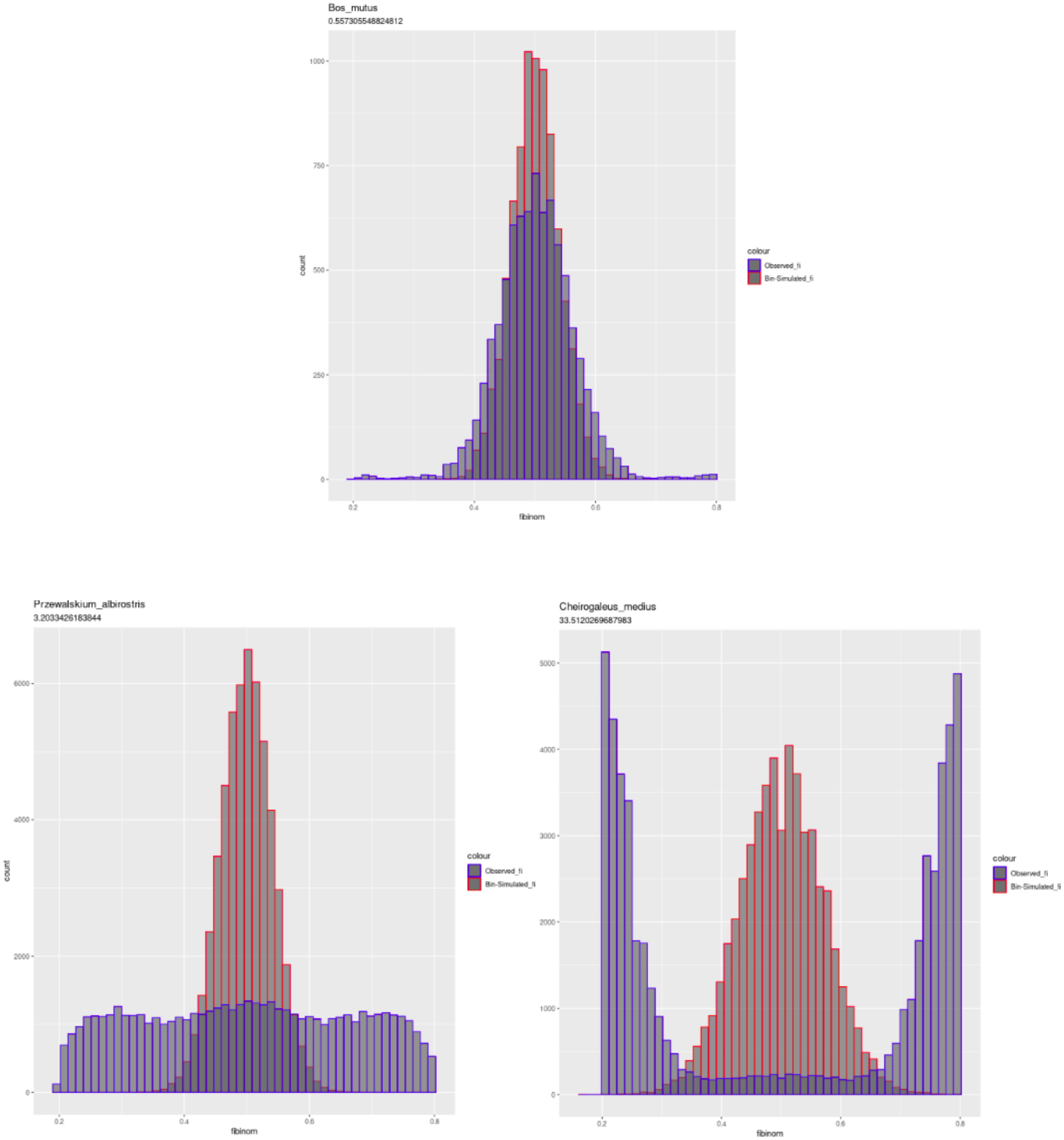


Figure S4: **Allelic balance distribution from three example**. In blue, distribution of locus frequencies (fi) for one species (*Bos mutus*, top) corresponding to the unimodal centered around 0.5 and two outliers species (*Cheirogaleus medius*, bottom right and *Przewalskium albirostris*, bottom left). The red histograms correspond to what would be expected from a binomial sampling centred at 0.5.

#### 5.4 $d_N/d_S$ reconstruction by FastCoevol using the 144 species tree

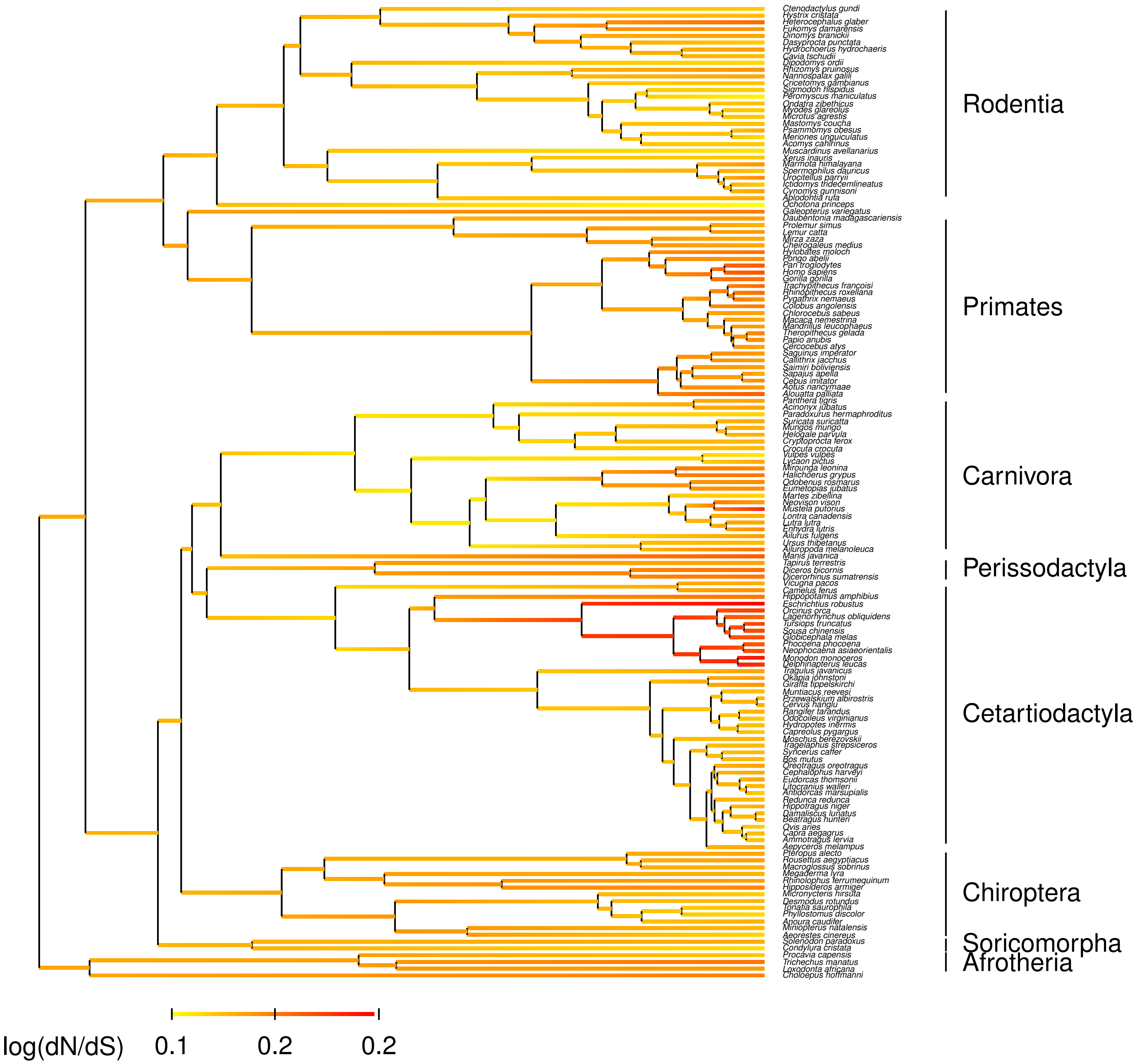


Figure S5: Reconstruction of $\log(d_N/d_S)$ along the reduced 144 mammalian species tree by FastCoevol

#### 5.5 Phylogenies used as input for FastCoevol

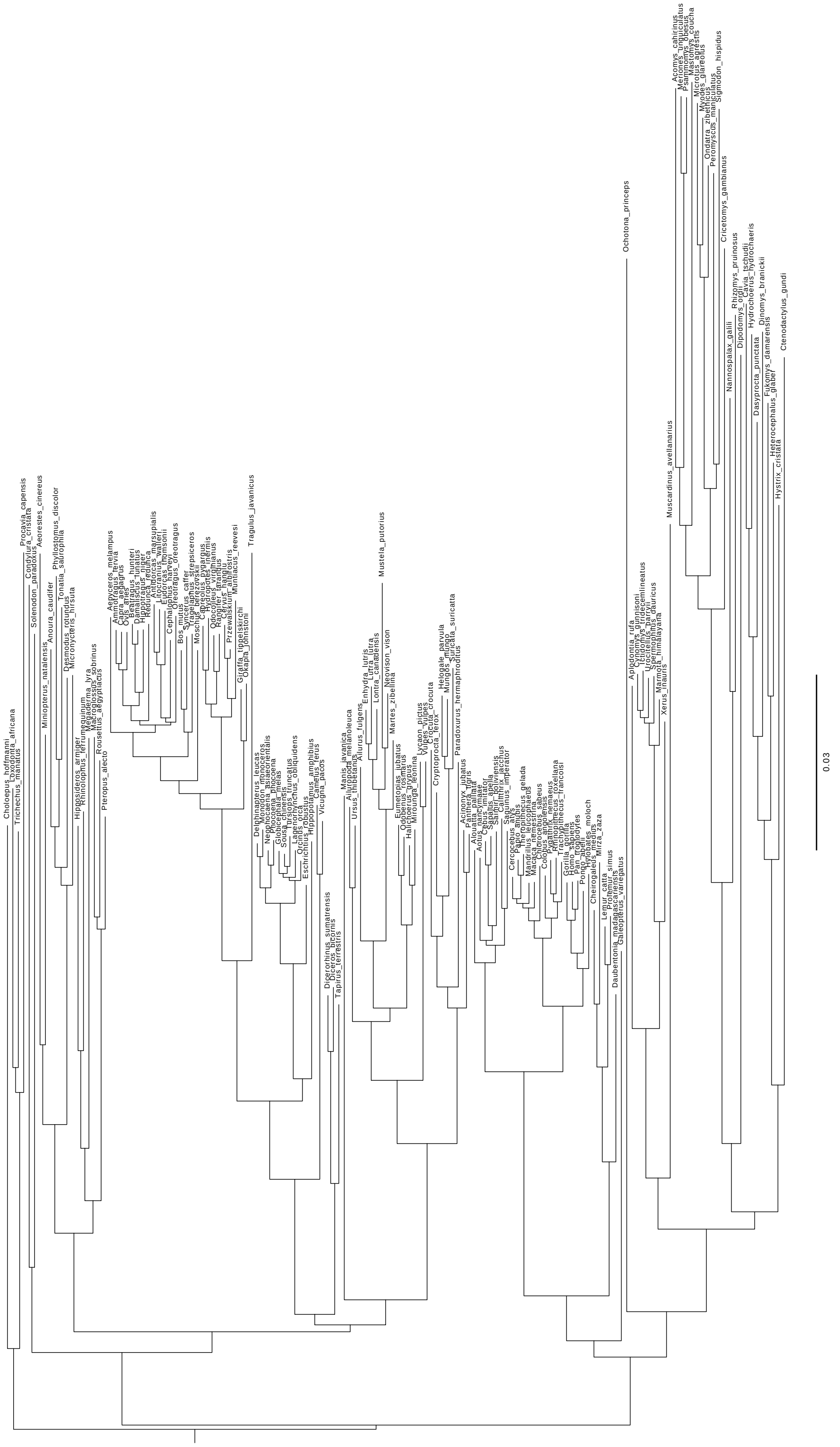


Figure S6: 144 mammals species phylogeny reconstructed using Iqtree

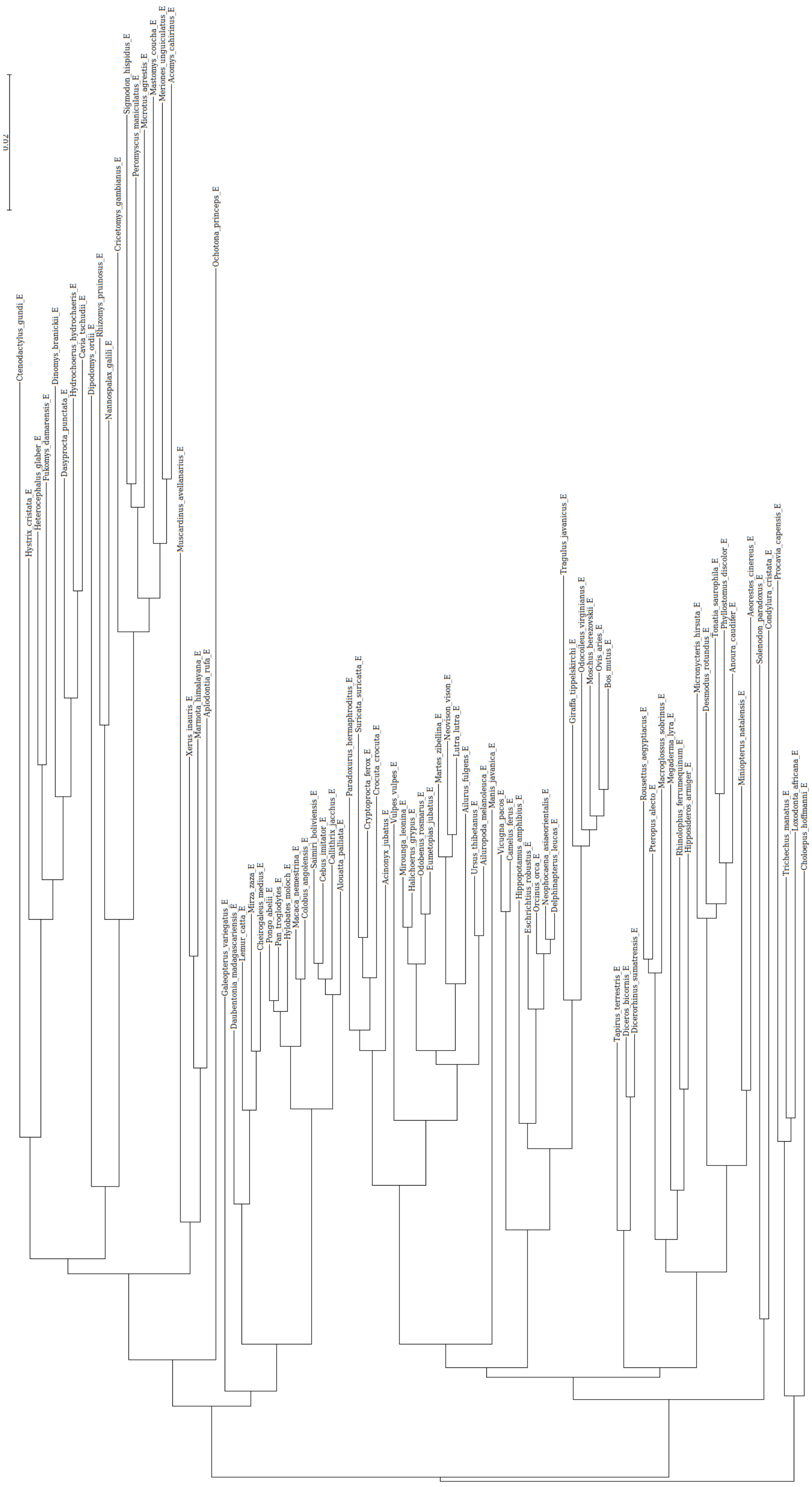


Figure S7: 89 mammals species phylogeny reconstructed using Iqtree

755 ## 5.6 Species sampling effect on the $d_S$ and $d_N/d_S$ estimation

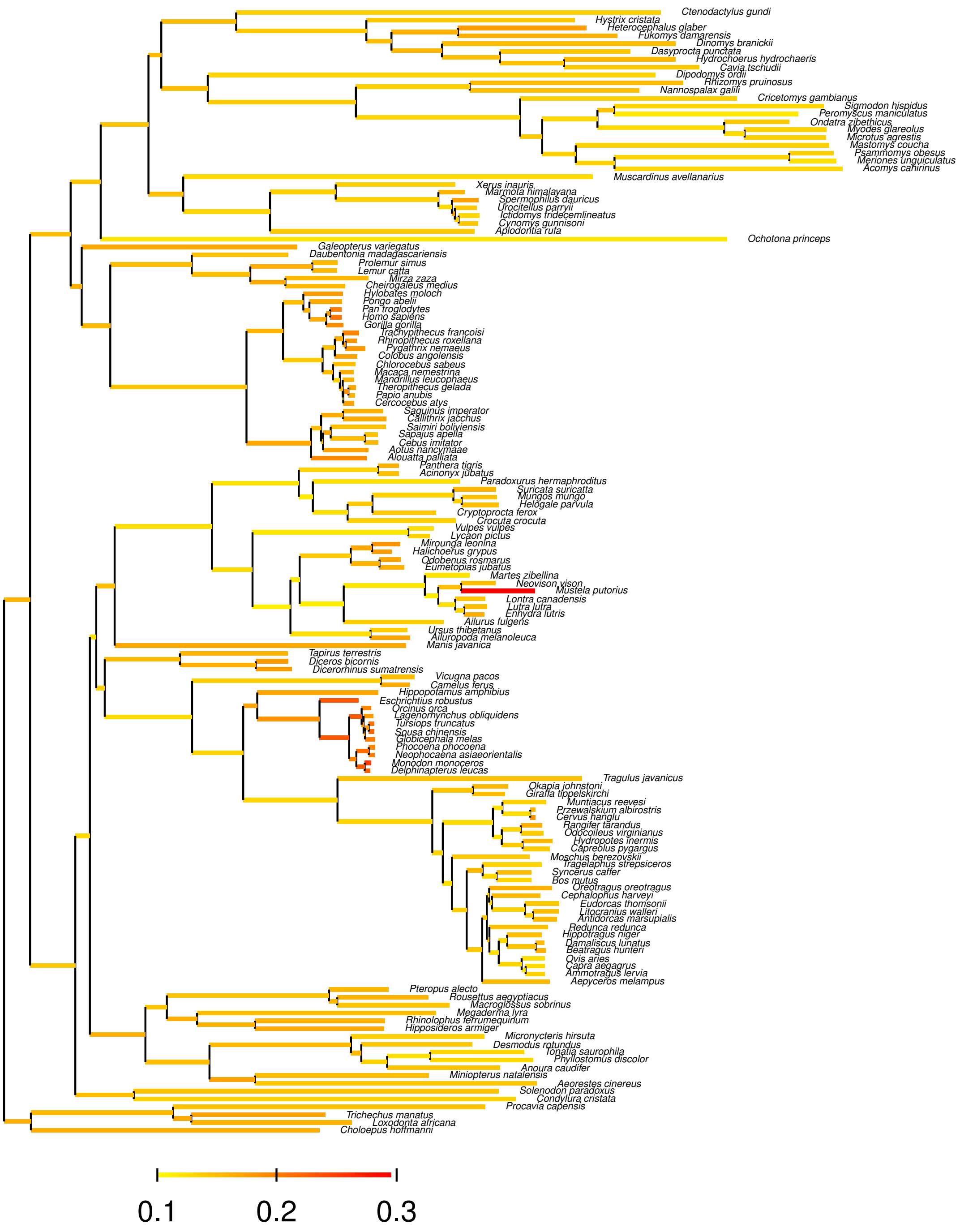


Figure S8: Empirical $d_S$-tree with branches colored as a function of empirical $d_N/d_S$ on the 144 species subset

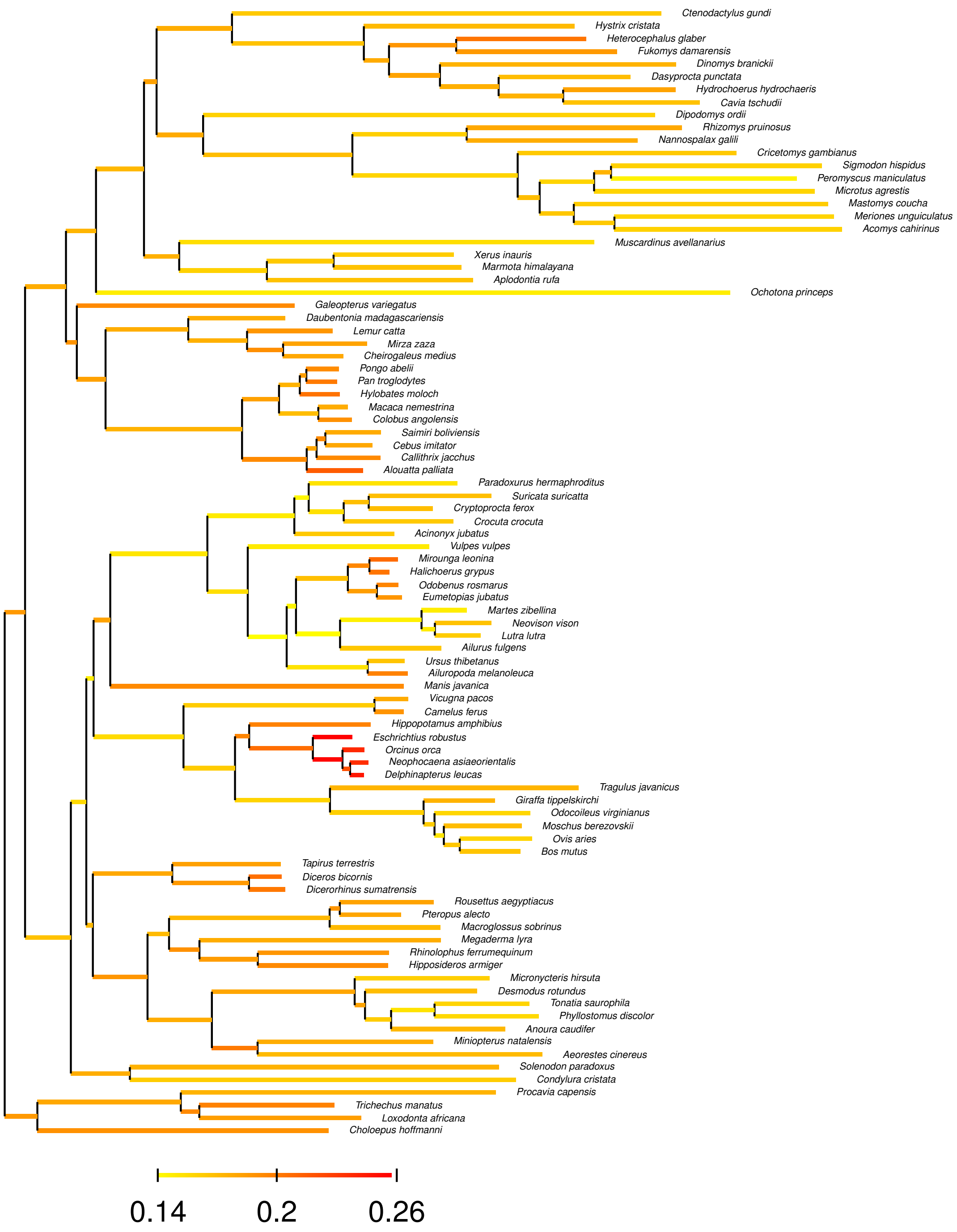


Figure S9: Empirical $d_S$-tree with branches colored as a function of empirical $d_N/d_S$ on the 89 species subset

##### 5.7 Decoupled estimation of $\pi_S$ and $\pi_N/\pi_S$

In our study, we estimate $\pi_S$ and reused it as denominator for $\pi_N/\pi_S$ estimation. This way to do has been controversial because this would mean studying the correlation between A and $\frac{B}{A}$, which are not independent measures, as it is suspected to create an artefactual negative correlation due to estimation errors on $\pi_S$.

We conducted a short analysis consisting in dividing randomly our 6002 genes list in two equal parts (named "1" and "2") and estimating $\pi_S$ only on one side and $\pi_N$ and $\pi_S$ to compute $\pi_N/\pi_S$ on the other side. This leads to a study of $A_1$ versus $\frac{B_2}{A_2}$ with both $A_i$ and $B_i$ corresponding to the same measure but on independent data. We did 500 times this random division of genes set and independant $\pi_S$ and $\pi_N/\pi_S$ computation in order to compute first, second and third quantile of our estimates.

We first compare graphically the relation between $\pi_N/\pi_S$ and the two type of $\pi_S$ estimations ($\frac{B_2}{A_2}$ versus $A_1$ or $A_2$). The linear regression didn't show strong differences at log scale (which is the scale of our study) with identical slope and correlation coefficient using the two types of $\pi_S$ measure (Figure S10). We then compare these slopes with the one obtained from the full dataset ($\frac{B}{A}$ versus A) used in the main results and didn't observe any difference.

Based on this analysis, we argue that using the same $\pi_S$ for both $\pi_N/\pi_S$ and $\pi_S$ measure isn't a problem because of the high quantity of SNPs used, as also previously noted (Leroy and Nabholz, 2022).

##### 5.8 Life history traits computation

In the article, we used the Anage database to obtain information on the mass, age of sexual maturity and longevity of each species considered in the study. To do so, we ordered the database by genus. For each genus, when there is more that one species represented, we compute the mean value for the trait over the genus and assign this mean as the observed trait value for the reference species. We use this method to minimize missing data. Thus, if a species under study is not present in the database, we can use information from a sister species. Here we quantify how much additional data this method has made it possible to add. As an example, concerning mass, 127 genus over the 144 of the analyses are represented in the database. 65 genus are represented once (53 by the species of the analysis) and 62 genus are represented by more than one species (50 including the species of the analysis). In the second case, we graphically explore the distribution of the species value compared to their genus mean (Figure S11). From this figure, we can conclude about a small intra-genus variance of the mass value, meaning that the use of a genus mean to represent the species add a small noise. However, this process allow adding 24 new mass measures. This conclusion is equivalent for maturity and longevity data.

##### 5.9 Robustness analysis

In the article, the removal of the polymorphism of a pre-determined set of six species with less than 1000 SNPs results in the observation of a significant negative relation between $\pi_S$ and mass, which is not significant when the polymorphism of these species is included. We thus question the sensibility of the analysis to the species sampling. To do so, we conducted different analyses using only the PGLS method to reduce the computational cost of the analysis.

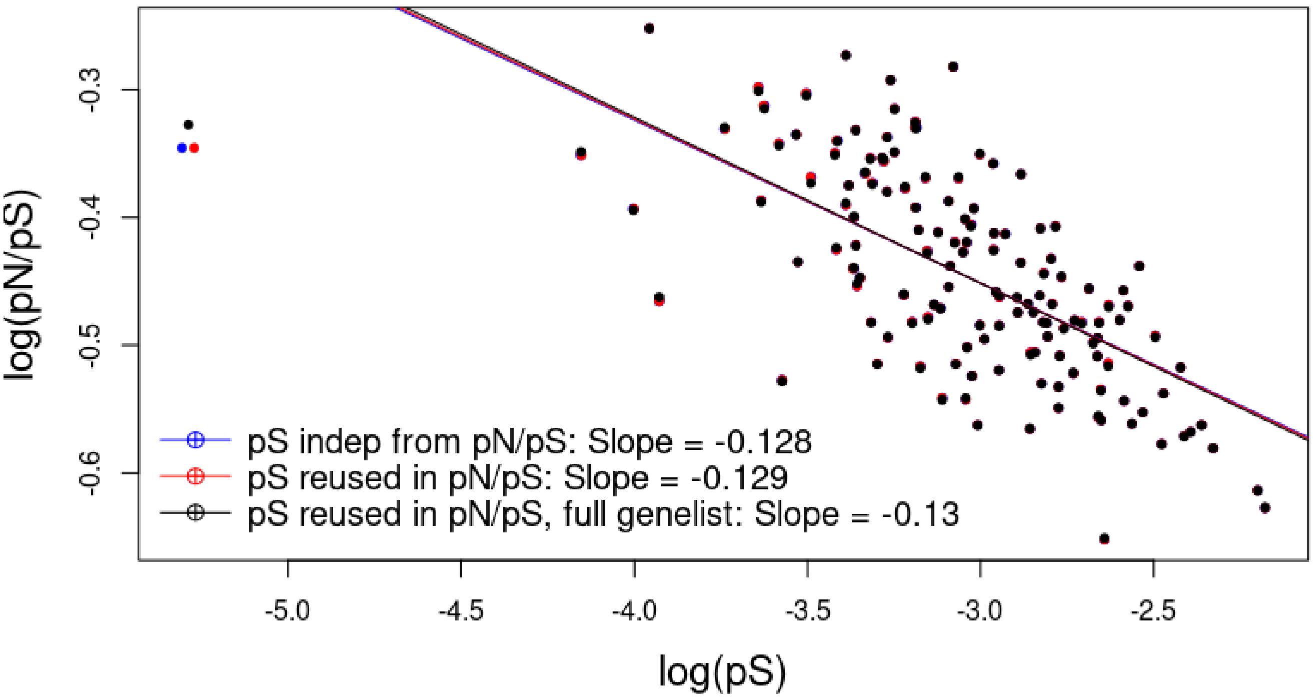


Figure S10: **Linear regression between log10($\pi_S$) and log10($\pi_N/\pi_S$).** The gene set is divided in two part to estimating independently two $\pi_S$, one for the $\pi_N/\pi_S$ estimation and the other for the $\pi_S$ values in itself. Blue points and slope correspond to a linear regression analysis between $\pi_S$ and $\pi_N/\pi_S$ with independent $\pi_S$ estimates for the two axes. Red points and slope correspond to a $\pi_S$ and $\pi_N/\pi_S$ comparison using twice the same $\pi_S$ estimates as in the main results (but with half of the gene set). Black points and slope correspond to a $\pi_S$ and $\pi_N/\pi_S$ comparison from the main results, with the full gene list.

First, we questioned the impact of removing the polymorphism of a seventh species and conducted a leave-one-out analysis, removing the polymorphism of each remaining species in turn. We did not observe any non-significant relation between $\pi_S$ and mass, nor between any of the other traits and a small variance for the $\pi_S$ mass correlation coefficient (0.00054). This suggests that the significant correlations observed are robust (in particular, they do not depend on a single data point)
Secondly, to test whether this observed correlation is due to the removal of polymorphism in these six particular species, we reran the analysis, randomly selecting six other species and removing their polymorphism instead of that in the six species used in the article. We repeated this analysis 50 times and failed to observe any $\pi_S$ mass correlation in 49 out of 50 cases, as was the case for the dataset containing all the polymorphism data. This suggest that this is the removal of the polymorphism of these six particular species that impact the significance of the correlation.

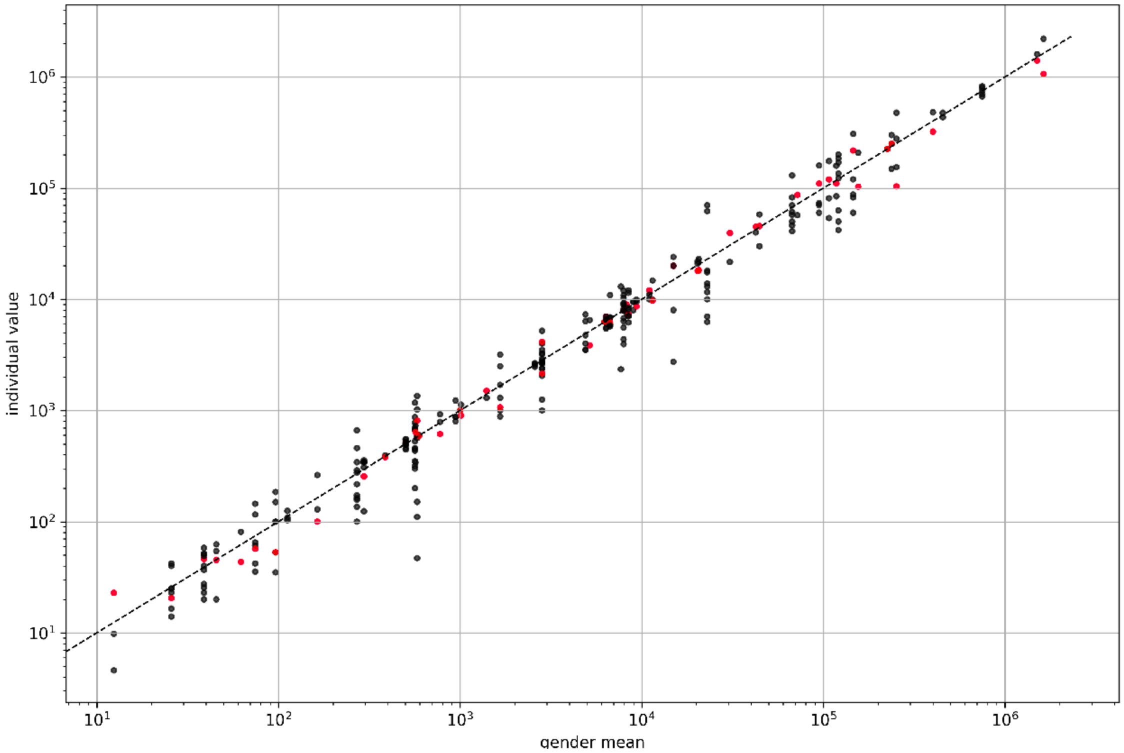


Figure S11: **Comparison of species mass values with mean mass values per gender**. Each column represents the genus variation (different genus can present a similar mean mass value and thus appear merged in the figure) and the red dots correspond to the direct mass value for the specie in the analysis. The dotted line represents y=x and illustrates the small variation in species values compared to the mean values for each gender.

Finally, we perform a linear regression analysis (without phylogenetic inertia correction) between $\pi_S$ and mass with three different subset : (1) with all the polymorphism data, (2) without the polymorphism of the six low $\pi_S$ species, (3) without the polymorphism of *Sigmodon hispidus*, one of the 6 low $\pi_S$ species. For the relation between log($\pi_S$) and log(mass), we don't observe any signifiant relation between log($\pi_S$) and log(mass). However, we can observe that *Sigmodon hispidus* alone has an important impact on the relation between $\pi_S$ and mass (respectively for the three subset: (1) slope=-0.26 and pval=0.399, (2) slope=-0.74 and pval=0.07, (3) slope=-0.6023 and pval=0.1).

These additional analyses show that it is definitely the six identified species that have an impact on the significance of the correlations. One of the species in this group (*Sigmodon Hispidus*) seems to have a particularly strong impact.

#### 5.10 Correlation matrix when include the six species with low polymorphism

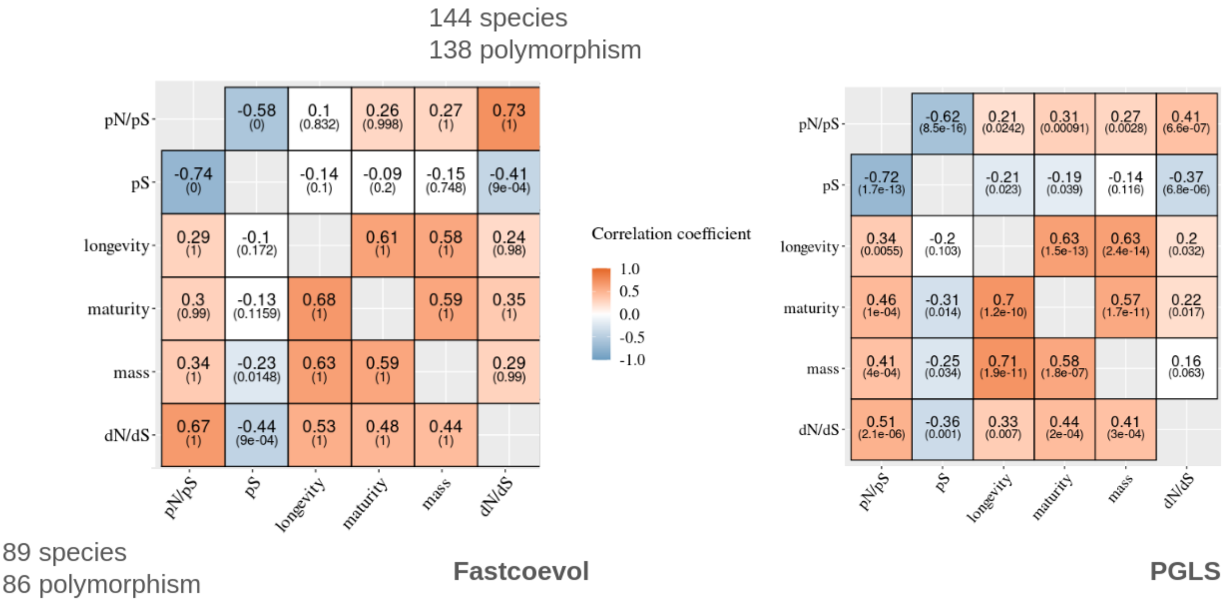


Figure S12: **Correlation coefficients from the FastCoevol and PGLS analyses**. **left**: FastCoevol analysis with the full polymorphism dataset. **right**: PGLS analysis with the full dataset. **Upper diagonal**: using the 144-species phylogeny. **Bottom diagonal**: using the 89-species phylogeny. Colours correspond to the magnitude of the correlation coefficients ($r$). Numbers in brackets correspond to the posterior probability for FastCoevol analyses (which must be higher than 0.975 or lower than 0.025 for the correlation to be considered significant) or p-value for PGLS analyses (which must be lower than 0.05 for the correlation to be considered as significant). Non-coloured estimates correspond to non-significant correlations.

##### 5.11 Summary tables

- **Per species genomic informations** : Table presenting for each species presente in the final dataset, the Busco score, the numer of final genes, the coverage, l50, n50 and presence/absence in the 89 species tree.

- Per species polymorphism informations : Table presenting for each species presente in the final dataset, the final number of SNPs, $\pi_S$, $\pi_N/\pi_S$, the number of genes with SNPs and the filtering of some species with abnormal VCF.

- **Per species summary of VCF filtering** : Table presenting for each species present in the final dataset, the number of SNPs after each VCF filtering steps.

- **Summary of the species dataset processing** : Table presenting for each species from the whole initial 197 species dataset, the different category of species filtering ending up with a set of 144 species. The two last column present the species present or not in the subdataset excluding six species with too low polymorphism and/or keeping only 10Ma separates species.

828

| sp | accesion_nb | Busco score | Nb finalgenes | Coverage | n50 | l50 | 10Matree |
|---|---|---|---|---|---|---|---|
| Acinonyx_jubatus | GCF_003709585.1 | 95.9 | 5877 | 105 | 48500042 | 15 | prst |
| Acomys_cahirinus | GCA_004027535.1 | 72.59 | 4739 | 29 | 65411 | 10134 | prst |
| Aeorestes_cinereus | GCA_011751065.1 | 90.17 | 5486 | 37 | 35075546 | 19 | prst |
| Aepyceros_melampus | GCA_006408695.1 | 74.57 | 4873 | 250 | 344542 | 2451 | none |
| Ailuropoda_melanoleuca | GCF_002007445.1 | 93.66 | 4955 | 108 | 129245720 | 8 | prst |
| Ailurus_fulgens | GCA_002007465.1 | 93.77 | 5760 | 117 | 2983736 | 215 | prst |
| Alouatta_palliata | GCA_004027835.1 | 70.44 | 4204 | 27 | 72427 | 11068 | prst |
| Ammotragus_lervia | GCA_002201775.1 | 93.8 | 5849 | 94 | 1301762 | 604 | none |
| Anoura_caudifer | GCA_004027475.1 | 87.94 | 5483 | 48 | 185021 | 3282 | prst |
| Antidorcas_marsupialis | GCA_006408585.1 | 82.27 | 5171 | 192 | 694905 | 1300 | none |
| Aotus_nancymaae | GCF_000952055.2 | 93.05 | 5747 | 189 | 8268663 | 102 | none |
| Aplodontia_rufa | GCA_004027875.1 | 72.22 | 4768 | 24 | 37811 | 19816 | prst |
| Beatragus_hunteri | GCA_004027495.1 | 75.27 | 4994 | 21 | 69303 | 11525 | none |
| Bos_mutus | GCA_007646595.3 | 93.07 | 5794 | 218 | 16589160 | 48 | prst |
| Callithrix_jacchus | GCA_009663435.2 | 92.99 | 5663 | 56 | 98198953 | 12 | prst |
| Camelus_ferus | GCF_009834535.1 | 96.03 | 5790 | 128 | 76025729 | 11 | prst |
| Capra_aegagrus | GCA_000765075.1 | 94.53 | 5882 | 172 | 1750063 | 409 | none |
| Capreolus_pygargus | GCA_012922965.1 | 92.81 | 5736 | 94 | 6067221 | 126 | none |
| Cavia_tschudii | GCA_004027695.1 | 84.25 | 5369 | 28 | 91436 | 8528 | prst |
| Cebus_imitator | GCF_001604975.1 | 93.11 | 5789 | 161 | 5274112 | 149 | prst |
| Cephalophus_harveyi | GCA_006410635.1 | 85.94 | 5479 | 197 | 365466 | 2293 | none |
| Cercocebus_atys | GCF_000955945.1 | 94.15 | 5817 | 17 | 12849131 | 66 | none |
| Cervus_hanglu | GCA_010411085.1 | 92.71 | 5717 | 80 | 77688133 | 13 | none |
| Cheirogaleus_medius | GCA_004024725.1 | 80.91 | 5228 | 67 | 118572 | 5555 | prst |
| Chlorocebus_sabeus | GCF_000409795.2 | 93.78 | 5768 | 197 | 81825804 | 14 | none |
| Choloepus_hoffmanni | GCA_000164785.2 | 84.34 | 5279 | 66 | 366442 | 2423 | prst |
| Colobus_angolensis | GCF_000951035.1 | 91.38 | 5708 | 150 | 7840981 | 117 | prst |
| Condylura_cristata | GCF_000260355.1 | 89.52 | 5520 | 195 | 55520359 | 13 | prst |
| Cricetomys_gambianus | GCA_004027575.1 | 83.94 | 5339 | 29 | 110049 | 5340 | prst |
| Crocuta_crocuta | GCA_008692635.1 | 94.63 | 5861 | 119 | 7236831 | 105 | prst |
| Cryptoprocta_ferox | GCA_004023885.1 | 86.52 | 5569 | 41 | 173473 | 3830 | prst |
| Ctenodactylus_gundi | GCA_004027205.1 | 91.89 | 5706 | 42 | 354548 | 1820 | prst |
| Cynomys_gunnisoni | GCA_011316645.1 | 91.95 | 5675 | 80 | 824613 | 911 | none |
| Damaliscus_lunatus | GCA_006408505.1 | 91.35 | 5692 | 163 | 1166796 | 799 | none |
| Dasyprocta_punctata | GCA_004363535.1 | 70.4 | 4559 | 34 | 43703 | 17174 | prst |
| Daubentonia_madagascariensis | GCA_004027145.1 | 92.42 | 5813 | 69 | 379919 | 1894 | prst |
| Delphinapterus_leucas | GCF_002288925.2 | 93.94 | 5841 | 106 | 31183418 | 21 | prst |
| Desmodus_rotundus | GCF_002940915.1 | 96.12 | 5827 | 123 | 26869735 | 26 | prst |
| Dicerorhinus_sumatrensis | GCA_002844835.1 | 93.85 | 5795 | 69 | 614498 | 1268 | prst |
| Diceros_bicornis | GCA_004027315.2 | 96.75 | 5885 | 10 | 14664201 | 54 | prst |
| Dinomys_branickii | GCA_004027595.1 | 80.54 | 5218 | 27 | 77918 | 9284 | prst |
| Dipodomys_ordii | GCF_000151885.1 | 89.71 | 5507 | 146 | 11931245 | 56 | prst |
| Enhydra_lutris | GCF_002288905.1 | 96.6 | 5884 | 55 | 38751465 | 20 | none |
| Eschrichtius_robustus | GCA_004363415.1 | 78.56 | 5200 | 34 | 94414 | 7799 | prst |
| Eudorcas_thomsonii | GCA_006408755.1 | 92.69 | 5374 | 220 | 1581717 | 484 | none |
| Eumetopias_jubatus | GCF_004028035.1 | 92.62 | 5744 | 43 | 14018600 | 54 | prst |
| Fukomys_damarensis | GCF_012274545.1 | 91.36 | 5571 | 172 | 62586000 | 15 | prst |
| Galeopterus_variegatus | GCA_004027255.2 | 94.59 | 5784 | 27 | 7885395 | 91 | prst |
| Giraffa_tippelskirchi | GCA_001651235.1 | 85.1 | 5469 | 37 | 212164 | 3486 | prst |
| Globicephala_melas | GCF_006547405.1 | 93.13 | 5832 | 48 | 18102937 | 42 | none |
| Gorilla_gorilla | GCF_008122165.1 | 91.79 | 5646 | 37 | 26116462 | 35 | none |
| Halichoerus_grypus | GCA_012393455.1 | 92.27 | 4443 | 26 | 1033864 | 637 | prst |

| | | | | | | | | |
|---|---|---|---|---|---|---|---|---|
| 829 | Helogale_parvula | GCA_004023845.1 | 83.72 | 5399 | 30 | 179119 | 3602 | none |
| | Heterocephalus_glaber | GCF_000247695.1 | 90.96 | 5552 | 150 | 20532749 | 42 | prst |
| | Hippopotamus_amphibius | GCA_004027065.2 | 93.56 | 3253 | 63 | 4444377 | 171 | prst |
| | Hipposideros_armiger | GCF_001890085.1 | 89.72 | 5552 | 186 | 2328177 | 272 | prst |
| | Hippotragus_niger | GCA_006942125.1 | 94.58 | 5867 | 58 | 4586323 | 168 | none |
| | Hydrochoerus_hydrochaeris | GCA_004027455.1 | 88.23 | 5514 | 25 | 202224 | 3717 | prst |
| | Hydropotes_inermis | GCA_006459105.1 | 94.18 | 5773 | 21 | 13818975 | 55 | none |
| | Hylobates_moloch | GCF_009828535.2 | 92.33 | 5617 | 64 | 125196221 | 9 | prst |
| | Hystrix_cristata | GCA_004026905.1 | 79.49 | 5105 | 29 | 64768 | 10348 | prst |
| | Ictidomys_tridecemlineatus | GCF_000236235.1 | 94.29 | 5692 | 64 | 8192786 | 80 | none |
| | Lagenorhynchus_obliquidens | GCF_003676395.1 | 93.8 | 5845 | 52 | 28371583 | 22 | none |
| | Lemur_catta | GCA_004024665.1 | 87.31 | 5509 | 66 | 215715 | 2945 | prst |
| | Litocranius_walleri | GCA_006410535.1 | 91.79 | 5627 | 232 | 3126223 | 282 | none |
| | Lontra_canadensis | GCF_010015895.1 | 95.53 | 5855 | 50 | 18460785 | 39 | none |
| | Loxodonta_africana | GCF_000001905.1 | 93.03 | 5732 | 42 | 46401353 | 21 | prst |
| | Lutra_lutra | GCA_902655055.1 | 95.56 | 5787 | 86 | 149004807 | 7 | prst |
| | Lycaon_pictus | GCA_004216515.1 | 84.74 | 4993 | 45 | 7494581 | 76 | none |
| | Macaca_nemestrina | GCF_000956065.1 | 94.0 | 5839 | 242 | 15219753 | 62 | prst |
| | Macroglossus_sobrinus | GCA_004027375.1 | 91.85 | 5709 | 48 | 453401 | 1154 | prst |
| | Mandrillus_leucophaeus | GCF_000951045.1 | 91.11 | 5720 | 104 | 3186748 | 285 | none |
| | Manis_javanica | GCF_001685135.1 | 85.45 | 5464 | 94 | 204728 | 3399 | prst |
| | Marmota_himalayana | GCA_005280165.1 | 91.47 | 5614 | 183 | 1497034 | 483 | prst |
| | Martes_zibellina | GCA_012583365.1 | 95.0 | 5822 | 104 | 5199373 | 134 | prst |
| | Mastomys_coucha | GCF_008632895.1 | 95.11 | 5706 | 56 | 118621203 | 8 | prst |
| | Megaderma_lyra | GCA_004026885.1 | 88.14 | 5554 | 38 | 96489 | 6165 | prst |
| | Meriones_unguiculatus | GCA_004026785.1 | 87.69 | 5581 | 28 | 100883 | 5504 | prst |
| | Micronycteris_hirsuta | GCA_004026765.1 | 78.83 | 5104 | 35 | 68868 | 8795 | prst |
| | Microtus_agrestis | GCA_902806755.1 | 89.98 | 5654 | 28 | 13349786 | 45 | prst |
| | Miniopterus_natalensis | GCF_001595765.1 | 93.83 | 5733 | 115 | 4315193 | 118 | prst |
| | Mirounga_leonina | GCF_011800145.1 | 93.62 | 5769 | 46 | 54232831 | 16 | prst |
| | Mirza_zaza | GCA_008750895.1 | 72.62 | 4752 | 56 | 75843 | 8667 | prst |
| | Monodon_monoceros | GCF_005190385.1 | 93.77 | 5772 | 42 | 107566389 | 9 | none |
| | Moschus_berezovskii | GCA_006459085.1 | 91.82 | 5169 | 272 | 2509225 | 326 | prst |
| | Mungos_mungo | GCA_004023785.1 | 88.37 | 5626 | 43 | 236501 | 2886 | none |
| | Muntiacus_reevesi | GCA_008787405.2 | 93.81 | 5723 | 19 | 94101870 | 7 | none |
| | Muscardinus_avellanarius | GCA_004027005.1 | 71.58 | 4712 | 29 | 59013 | 12037 | prst |
| | Mustela_putorius | GCA_009859225.1 | 72.59 | 4568 | 157 | 33389539 | 24 | none |
| | Myodes_glareolus | GCA_004368595.1 | 78.34 | 4760 | 195 | 1590265 | 413 | none |
| | Nannospalax_galili | GCF_000622305.1 | 94.89 | 5809 | 62 | 3618479 | 238 | prst |
| | Neophocaena_asiaeorientalis | GCF_003031525.1 | 93.45 | 5855 | 251 | 6341296 | 103 | prst |
| | Neovison_vison | GCA_900108605.1 | 94.34 | 5320 | 64 | 6814223 | 103 | prst |
| | Ochotona_princeps | GCF_000292845.1 | 91.43 | 5639 | 136 | 26863993 | 26 | prst |
| | Odobenus_rosmarus | GCF_000321225.1 | 93.96 | 5818 | 35 | 2616778 | 269 | prst |
| | Odocoileus_virginianus | GCF_002102435.1 | 91.35 | 5747 | 233 | 850721 | 758 | prst |
| | Okapia_johnstoni | GCA_001660835.1 | 81.69 | 5302 | 32 | 111538 | 6512 | none |
| | Ondatra_zibethicus | GCA_004026605.1 | 81.71 | 5304 | 39 | 89093 | 7754 | none |
| | Orcinus_orca | GCF_000331955.2 | 93.98 | 5861 | 216 | 12735091 | 60 | prst |
| | Oreotragus_oreotragus | GCA_006410675.1 | 79.14 | 4750 | 152 | 339390 | 3115 | none |
| | Ovis_aries | GCA_000765115.1 | 94.1 | 5869 | 174 | 2217029 | 328 | prst |
| | Pan_troglodytes | GCF_002880755.1 | 94.23 | 5787 | 71 | 53103722 | 19 | prst |
| | Panthera_tigris | GCF_000464555.1 | 86.47 | 5477 | 131 | 8860407 | 87 | none |
| | Papio_anubis | GCF_008728515.1 | 92.02 | 5654 | 54 | 140274886 | 9 | none |
| | Paradoxurus_hermaphroditus | GCA_004024585.1 | 74.2 | 4833 | 32 | 71823 | 8704 | prst |

| | | | | | | | | |
|---|---|---|---|---|---|---|---|---|
| 830 | Peromyscus_maniculatus | GCF_000500345.1 | 94.88 | 5813 | 110 | 3760915 | 193 | prst |
| | Phocoena_phocoena | GCA_004363495.1 | 82.16 | 5345 | 47 | 115969 | 6441 | none |
| | Phyllostomus_discolor | GCF_004126475.1 | 96.15 | 5762 | 98 | 110241909 | 6 | prst |
| | Pongo_abelii | GCF_002880775.1 | 94.97 | 5801 | 41 | 98475126 | 13 | prst |
| | Procavia_capensis | GCA_004026925.2 | 94.7 | 5814 | 19 | 9071062 | 110 | prst |
| | Prolemur_simus | GCA_003258685.1 | 94.91 | 5837 | 350 | 2710671 | 251 | none |
| | Przewalskium_albirostris | GCA_006408465.1 | 89.79 | 5653 | 203 | 3769372 | 218 | none |
| | Psammomys_obesus | GCA_002215935.2 | 95.83 | 5872 | 154 | 10472398 | 62 | none |
| | Pteropus_alecto | GCF_000325575.1 | 95.59 | 5849 | 158 | 15954802 | 36 | prst |
| | Pygathrix_nemaeus | GCA_004024825.1 | 73.31 | 4823 | 46 | 68569 | 12661 | none |
| | Rangifer_tarandus | GCA_004026565.1 | 80.1 | 5259 | 39 | 89062 | 8918 | none |
| | Redunca_redunca | GCA_006410935.1 | 71.3 | 4498 | 238 | 423407 | 1625 | none |
| | Rhinolophus_ferrumequinum | GCF_004115265.1 | 96.13 | 5859 | 78 | 88025743 | 11 | prst |
| | Rhinopithecus_roxellana | GCF_007565055.1 | 93.19 | 5714 | 100 | 144559847 | 9 | none |
| | Rhizomys_pruinosus | GCA_009823505.1 | 89.82 | 5575 | 16 | 2203772 | 489 | prst |
| | Rousettus_aegyptiacus | GCF_001466805.2 | 95.89 | 5826 | 50 | 2007187 | 297 | prst |
| | Saguinus_imperator | GCA_004024885.1 | 76.11 | 5023 | 44 | 65636 | 12800 | none |
| | Saimiri_boliviensis | GCF_000235385.1 | 92.62 | 5751 | 158 | 18744880 | 39 | prst |
| | Sapajus_apella | GCF_009761245.1 | 92.71 | 5759 | 34 | 23742480 | 35 | none |
| | Sigmodon_hispidus | GCA_004025045.1 | 82.44 | 5307 | 38 | 101373 | 7354 | prst |
| | Solenodon_paradoxus | GCA_004363575.1 | 91.26 | 5692 | 27 | 407682 | 1301 | prst |
| | Sousa_chinensis | GCA_007760645.1 | 93.44 | 5825 | 148 | 19436979 | 39 | none |
| | Spermophilus_dauricus | GCA_002406435.1 | 88.97 | 5481 | 233 | 1761345 | 493 | none |
| | Suricata_suricatta | GCF_006229205.1 | 93.85 | 5704 | 44 | 141453419 | 8 | prst |
| | Syncerus_caffer | GCA_902825105.1 | 94.44 | 5804 | 198 | 69160875 | 13 | none |
| | Tapirus_terrestris | GCA_004025025.1 | 84.83 | 5379 | 38 | 186384 | 3791 | prst |
| | Theropithecus_gelada | GCF_003255815.1 | 93.5 | 5744 | 76 | 130230028 | 9 | none |
| | Tonatia_saurophila | GCA_004024845.1 | 86.02 | 5410 | 42 | 165561 | 3525 | prst |
| | Trachypithecus_francoisi | GCF_009764315.1 | 93.29 | 5746 | 39 | 130977661 | 10 | none |
| | Tragelaphus_strepsiceros | GCA_006410795.1 | 84.0 | 5423 | 241 | 511483 | 1640 | none |
| | Tragulus_javanicus | GCA_004024965.2 | 93.43 | 5730 | 36 | 14082842 | 49 | prst |
| | Trichechus_manatus | GCF_000243295.1 | 94.26 | 5781 | 187 | 14442683 | 67 | prst |
| | Tursiops_truncatus | GCF_011762595.1 | 93.3 | 5776 | 352 | 108430135 | 9 | none |
| | Urocitellus_parryii | GCF_003426925.1 | 94.49 | 5752 | 122 | 3964291 | 175 | none |
| | Ursus_thibetanus | GCA_009660055.1 | 95.24 | 5850 | 98 | 26803000 | 27 | prst |
| | Vicugna_pacos | GCA_000767525.1 | 95.06 | 5797 | 137 | 5303709 | 107 | prst |
| | Vulpes_vulpes | GCF_003160815.1 | 94.16 | 5771 | 145 | 12472085 | 55 | prst |
| | Xerus_inauris | GCA_004024805.1 | 77.21 | 5064 | 21 | 83865 | 8399 | prst |

| 831 | sp | nb_snp | pS | pN/pS | nbgeneswithsnp | vcf_type |
|---|---|---|---|---|---|---|
| | Acinonyx_jubatus | 3025 | 0.00044 | 0.37834 | 1789 | normal |
| | Acomys_cahirinus | 1114 | NA | NA | 28 | abnormal vcf |
| | Aeorestes_cinereus | 34281 | 0.00663 | 0.23602 | 4518 | normal |
| | Aepyceros_melampus | 4815 | 0.00113 | 0.34563 | 1972 | normal |
| | Ailuropoda_melanoleuca | 3966 | 0.00091 | 0.38094 | 1769 | normal |
| | Ailurus_fulgens | 6003 | 0.00096 | 0.40461 | 2570 | normal |
| | Alouatta_palliata | 732 | 0.00018 | 0.46825 | 410 | too few snp |
| | Ammotragus_lervia | 6630 | 0.00103 | 0.31973 | 2208 | normal |
| | Anoura_caudifer | 8364 | 0.0015 | 0.29506 | 3054 | normal |
| | Antidorcas_marsupialis | 17281 | 0.00404 | 0.27044 | 4096 | normal |
| | Aotus_nancymaae | 18646 | 0.00287 | 0.3646 | 4436 | normal |
| | Aplodontia_rufa | 4514 | 0.0009 | 0.39706 | 2063 | normal |
| | Beatragus_hunteri | 583 | 0.00012 | 0.34493 | 427 | too few snp |
| | Bos_mutus | 8208 | 0.00127 | 0.34479 | 3183 | normal |
| | Callithrix_jacchus | 6405 | 0.00094 | 0.39181 | 2492 | normal |
| | Camelus_ferus | 5155 | 0.00082 | 0.36464 | 2264 | normal |
| | Capra_aegagrus | 9007 | 0.00137 | 0.34068 | 3224 | normal |
| | Capreolus_pygargus | 18079 | 0.00337 | 0.28982 | 4489 | normal |
| | Cavia_tschudii | 2210 | 0.00038 | 0.37698 | 981 | normal |
| | Cebus_imitator | 3999 | 0.00054 | 0.45991 | 1905 | normal |
| | Cephalophus_harveyi | 8962 | NA | NA | 3011 | abnormal vcf |
| | Cercocebus_atys | 4280 | 0.00071 | 0.33139 | 2270 | normal |
| | Cervus_hanglu | 9298 | 0.00148 | 0.34573 | 2795 | normal |
| | Cheirogaleus_medius | 52463 | NA | NA | 3546 | abnormal vcf |
| | Chlorocebus_sabeus | 7449 | 0.00109 | 0.387 | 2586 | normal |
| | Choloepus_hoffmanni | 4877 | 0.00086 | 0.42825 | 1682 | normal |
| | Colobus_angolensis | 7013 | 0.00099 | 0.44668 | 2912 | normal |
| | Condylura_cristata | 9197 | 0.00168 | 0.28259 | 3260 | normal |
| | Cricetomys_gambianus | 4660 | 0.00091 | 0.28743 | 2188 | normal |
| | Crocuta_crocuta | 6059 | 0.00099 | 0.32773 | 2582 | normal |
| | Cryptoprocta_ferox | 4323 | 0.00077 | 0.28657 | 2393 | normal |
| | Ctenodactylus_gundi | 6818 | 0.00111 | 0.34801 | 2913 | normal |
| | Cynomys_gunnisoni | 2189 | 0.00032 | 0.42363 | 740 | normal |
| | Damaliscus_lunatus | 7830 | 0.00139 | 0.31089 | 3073 | normal |
| | Dasyprocta_punctata | 12324 | 0.00294 | 0.2802 | 2967 | normal |
| | Daubentonia_madagascariensis | 3241 | 0.00044 | 0.46573 | 1834 | normal |
| | Delphinapterus_leucas | 4387 | 0.0006 | 0.42086 | 2400 | normal |
| | Desmodus_rotundus | 25734 | 0.00435 | 0.27372 | 4740 | normal |
| | Dicerorhinus_sumatrensis | 5587 | 0.00081 | 0.40993 | 2261 | normal |
| | Diceros_bicornis | 627 | 0.0001 | 0.40346 | 523 | too few snp |
| | Dinomys_branickii | 1394 | 0.00023 | 0.5 | 743 | normal |
| | Dipodomys_ordii | 2702 | 0.00048 | 0.32934 | 1419 | normal |
| | Enhydra_lutris | 1657 | 0.00023 | 0.40944 | 1155 | normal |
| | Eschrichtius_robustus | 2010 | 0.00031 | 0.49604 | 1404 | normal |
| | Eudorcas_thomsonii | 7486 | 0.00165 | 0.3914 | 2439 | normal |
| | Eumetopias_jubatus | 1978 | 0.0003 | 0.36754 | 1238 | normal |
| | Fukomys_damarensis | 3862 | 0.00055 | 0.50994 | 1875 | normal |
| | Galeopterus_variegatus | 13602 | 0.00218 | 0.32029 | 3350 | normal |
| | Giraffa_tippelskirchi | 2584 | 0.00041 | 0.40858 | 1534 | normal |
| | Globicephala_melas | 3475 | 0.00049 | 0.42273 | 1775 | normal |
| | Gorilla_gorilla | 7285 | 0.00109 | 0.43879 | 2916 | normal |
| | Halichoerus_grypus | 1437 | 0.00045 | 0.3574 | 807 | normal |

| | | | | | | |
|---|---|---|---|---|---|---|
| 832 | Helogale_parvula | 2977 | 0.00054 | 0.32101 | 1321 | normal |
| | Heterocephalus_glaber | 3087 | 0.00041 | 0.53313 | 1403 | normal |
| | Hippopotamus_amphibius | 4762 | 0.00171 | 0.35802 | 1642 | normal |
| | Hipposideros_armiger | 9633 | 0.00161 | 0.34028 | 2886 | normal |
| | Hippotragus_niger | 7609 | 0.00128 | 0.33542 | 2993 | normal |
| | Hydrochoerus_hydrochaeris | 2928 | 0.00046 | 0.43189 | 1863 | normal |
| | Hydropotes_inermis | 3209 | 0.00067 | 0.30346 | 1790 | normal |
| | Hylobates_moloch | 8945 | 0.00131 | 0.43041 | 3391 | normal |
| | Hystrix_cristata | 10945 | 0.0022 | 0.32904 | 3537 | normal |
| | Ictidomys_tridecemlineatus | 7933 | 0.00142 | 0.33558 | 2361 | normal |
| | Lagenorhynchus_obliquidens | 5195 | 0.00075 | 0.38743 | 2488 | normal |
| | Lemur_catta | 11048 | 0.00187 | 0.33055 | 3296 | normal |
| | Litocranius_walleri | 2476 | NA | NA | 1248 | abnormal vcf |
| | Lontra_canadensis | 4998 | 0.00077 | 0.33775 | 2148 | normal |
| | Loxodonta_africana | 3685 | 0.00053 | 0.44197 | 1791 | normal |
| | Lutra_lutra | 2741 | 0.00043 | 0.36358 | 1187 | normal |
| | Lycaon_pictus | 2053 | 0.00039 | 0.45666 | 1070 | normal |
| | Macaca_nemestrina | 15490 | 0.00234 | 0.33893 | 4467 | normal |
| | Macroglossus_sobrinus | 12011 | 0.00196 | 0.32869 | 3273 | normal |
| | Mandrillus_leucophaeus | 8493 | 0.00131 | 0.36681 | 3101 | normal |
| | Manis_javanica | 9512 | 0.0016 | 0.36931 | 3159 | normal |
| | Marmota_himalayana | 2784 | 0.00042 | 0.42185 | 1619 | normal |
| | Martes_zibellina | 10149 | 0.00168 | 0.29351 | 3194 | normal |
| | Mastomys_coucha | 1708 | NA | NA | 181 | abnormal vcf |
| | Megaderma_lyra | 5321 | 0.00091 | 0.31485 | 2555 | normal |
| | Meriones_unguiculatus | 1489 | 0.00027 | 0.29626 | 599 | normal |
| | Micronycteris_hirsuta | 16030 | 0.00333 | 0.2645 | 3557 | normal |
| | Microtus_agrestis | 12976 | 0.00234 | 0.30446 | 3527 | normal |
| | Miniopterus_natalensis | 4841 | 0.00085 | 0.30549 | 2364 | normal |
| | Mirounga_leonina | 8972 | 0.00144 | 0.31195 | 3429 | normal |
| | Mirza_zaza | 5285 | 0.00113 | 0.32733 | 1873 | normal |
| | Monodon_monoceros | 1982 | 0.00029 | 0.46214 | 1429 | normal |
| | Moschus_berezovskii | 9356 | 0.00267 | 0.3391 | 3140 | normal |
| | Mungos_mungo | 9116 | 0.00152 | 0.32948 | 3244 | normal |
| | Muntiacus_reevesi | 3848 | 0.00063 | 0.32895 | 1292 | normal |
| | Muscardinus_avellanarius | 366 | 7E-05 | 0.44819 | 200 | too few snp |
| | Mustela_putorius | 1203 | 0.00024 | 0.48445 | 537 | normal |
| | Myodes_glareolus | 8242 | 0.00169 | 0.3101 | 2063 | normal |
| | Nannospalax_galili | 4842 | 0.00081 | 0.35127 | 2328 | normal |
| | Neophocaena_asiaeorientalis | 4578 | 0.0007 | 0.37333 | 2200 | normal |
| | Neovison_vison | 3381 | 0.00073 | 0.34021 | 1578 | normal |
| | Ochotona_princeps | 9286 | 0.00155 | 0.32911 | 3207 | normal |
| | Odobenus_rosmarus | 3954 | 0.0006 | 0.34668 | 2285 | normal |
| | Odocoileus_virginianus | 21995 | 0.00387 | 0.2682 | 3717 | normal |
| | Okapia_johnstoni | 3827 | 0.00066 | 0.38917 | 2139 | normal |
| | Ondatra_zibethicus | 4958 | 0.00094 | 0.29916 | 1746 | normal |
| | Orcinus_orca | 2780 | 0.00038 | 0.44558 | 1522 | normal |
| | Oreotragus_oreotragus | 10688 | 0.00258 | 0.34906 | 3038 | normal |
| | Ovis_aries | 15970 | 0.0026 | 0.28616 | 3750 | normal |
| | Pan_troglodytes | 4323 | 0.00056 | 0.48384 | 2338 | normal |
| | Panthera_tigris | 2656 | 0.00044 | 0.35347 | 1295 | normal |
| | Papio_anubis | 2880 | 0.00043 | 0.39888 | 1638 | normal |
| | Paradoxurus_hermaphroditus | 10174 | 0.00219 | 0.2783 | 2943 | normal |

| | | | | | | |
|---|---|---|---|---|---|---|
| 833 | Peromyscus_maniculatus | 12272 | 0.00228 | 0.22333 | 1372 | normal |
| | Phocoena_phocoena | 6260 | 0.00109 | 0.37503 | 2858 | normal |
| | Phyllostomus_discolor | 34619 | 0.00631 | 0.24334 | 4797 | normal |
| | Pongo_abelii | 10236 | 0.00149 | 0.39024 | 3387 | normal |
| | Procavia_capensis | 15909 | 0.00274 | 0.27454 | 4318 | normal |
| | Prolemur_simus | 13305 | 0.00223 | 0.27616 | 3863 | normal |
| | Przewalskium_albirostris | 54210 | NA | NA | 5475 | abnormal vcf |
| | Psammomys_obesus | 3923 | 0.00056 | 0.44804 | 1771 | normal |
| | Pteropus_alecto | 26581 | 0.00469 | 0.26268 | 4852 | normal |
| | Pygathrix_nemaeus | 4686 | 0.00089 | 0.37393 | 2117 | normal |
| | Rangifer_tarandus | 8520 | 0.00156 | 0.32128 | 3118 | normal |
| | Redunca_redunca | 7403 | 0.00174 | 0.32567 | 2556 | normal |
| | Rhinolophus_ferrumequinum | 8204 | 0.00139 | 0.27207 | 2519 | normal |
| | Rhinopithecus_roxellana | 1893 | 0.00026 | 0.45317 | 1126 | normal |
| | Rhizomys_pruinosus | 4347 | 0.00065 | 0.46729 | 1857 | normal |
| | Rousettus_aegyptiacus | 11282 | 0.00185 | 0.30082 | 2681 | normal |
| | Saguinus_imperator | 6623 | 0.00118 | 0.38638 | 2543 | normal |
| | Saimiri_boliviensis | 4439 | 0.00065 | 0.40518 | 1802 | normal |
| | Sapajus_apella | 5700 | 0.00084 | 0.38017 | 2169 | normal |
| | Sigmodon_hispidus | 33 | 1E-05 | 0.47059 | 11 | too few snp |
| | Solenodon_paradoxus | 3513 | 0.00054 | 0.41675 | 1840 | normal |
| | Sousa_chinensis | 888 | 0.00011 | 0.55967 | 551 | too few snp |
| | Spermophilus_dauricus | 5946 | 0.00113 | 0.30221 | 2495 | normal |
| | Suricata_suricatta | 15597 | 0.00252 | 0.33095 | 4009 | normal |
| | Syncerus_caffer | 19532 | 0.0032 | 0.32081 | 4234 | normal |
| | Tapirus_terrestris | 11797 | 0.00206 | 0.3503 | 3667 | normal |
| | Theropithecus_gelada | 3820 | 0.00052 | 0.44321 | 1815 | normal |
| | Tonatia_saurophila | 12054 | 0.00223 | 0.29136 | 3415 | normal |
| | Trachypithecus_francoisi | 4952 | 0.00065 | 0.47212 | 2511 | normal |
| | Tragelaphus_strepsiceros | 7964 | 0.00152 | 0.35949 | 2904 | normal |
| | Tragulus_javanicus | 23253 | 0.00378 | 0.30377 | 4283 | normal |
| | Trichechus_manatus | 4850 | 0.00069 | 0.42841 | 1539 | normal |
| | Tursiops_truncatus | 3494 | 0.00048 | 0.44229 | 1626 | normal |
| | Urocitellus_parryii | 6172 | 0.00083 | 0.52253 | 1906 | normal |
| | Ursus_thibetanus | 13519 | 0.00217 | 0.31001 | 3724 | normal |
| | Vicugna_pacos | 11796 | 0.00211 | 0.31724 | 2905 | normal |
| | Vulpes_vulpes | 5541 | 0.00098 | 0.27367 | 2126 | normal |
| | Xerus_inauris | 2468 | 0.0005 | 0.30548 | 1292 | normal |

| | | | | | | | | | |
|---|---|---|---|---|---|---|---|---|---|
| 834 | species | raw_snp | biallelic_snp | call_snp | coding_snp | genelist_snp | hetero_snp | qual_snp | allelic_balance |
| | Choloepus_hoffmanni | 3867911 | 3867392 | 3401371 | 7445 | 5236 | 5222 | 4878 | 4877 |
| | Loxodonta_africana | 6348963 | 6343331 | 5438917 | 11318 | 7639 | 4154 | 3689 | 3685 |
| | Trichechus_manatus | 3114185 | 3112578 | 2724745 | 7784 | 5188 | 5132 | 4871 | 4850 |
| | Procavia_capensis | 16458011 | 16457830 | 16099833 | 32444 | 22215 | 22211 | 15919 | 15909 |
| | Condylura_cristata | 3461962 | 3459798 | 3199650 | 13629 | 9405 | 9390 | 9225 | 9197 |
| | Solenodon_paradoxus | 1687072 | 1686976 | 1561628 | 6435 | 4513 | 4513 | 3514 | 3513 |
| | Aeorestes_cinereus | 13235902 | 13235152 | 12737699 | 101003 | 72671 | 72643 | 34326 | 34281 |
| | Miniopterus_natalensis | 2249627 | 2249145 | 1989441 | 7515 | 4954 | 4928 | 4841 | 4841 |
| | Anoura_caudifer | 3813786 | 3813688 | 3680239 | 12369 | 8575 | 8575 | 8365 | 8364 |
| | Phyllostomus_discolor | 14766295 | 14765094 | 13909805 | 52198 | 35257 | 35049 | 34632 | 34619 |
| | Tonatia_saurophila | 5250643 | 5250543 | 5129431 | 17856 | 12547 | 12547 | 12054 | 12054 |
| | Desmodus_rotundus | 11742968 | 11738825 | 11182434 | 41464 | 27022 | 26028 | 25744 | 25734 |
| | Micronycteris_hirsuta | 8425912 | 8425801 | 8240018 | 22843 | 16953 | 16951 | 16037 | 16030 |
| | Hipposideros_armiger | 4436021 | 4434281 | 3943378 | 14878 | 10063 | 10026 | 9670 | 9633 |
| | Rhinolophus_ferrumequinum | 4562708 | 4562551 | 4372112 | 12991 | 8815 | 8751 | 8217 | 8204 |
| | Megaderma_lyra | 3680053 | 3679443 | 3115185 | 8266 | 5578 | 5578 | 5323 | 5321 |
| | Macroglossus_sobrinus | 3887865 | 3887737 | 3778439 | 18082 | 12292 | 12292 | 12016 | 12011 |
| | Rousettus_aegyptiacus | 13747426 | 13732212 | 13268215 | 53475 | 35118 | 11764 | 11290 | 11282 |
| | Pteropus_alecto | 12191444 | 12188675 | 11253289 | 41356 | 27426 | 27012 | 26601 | 26581 |
| | Aepyceros_melampus | 4256453 | 4254876 | 3443940 | 8655 | 6464 | 6430 | 4972 | 4815 |
| | Ammotragus_lervia | 4458698 | 4457469 | 3969977 | 10376 | 6853 | 6840 | 6643 | 6630 |
| | Capra_aegagrus | 5569313 | 5567129 | 5101678 | 14250 | 9313 | 9305 | 9057 | 9007 |
| | Ovis_aries | 9419714 | 9416861 | 8827788 | 24397 | 16269 | 16244 | 16043 | 15970 |
| | Beatragus_hunteri | 955550 | 955370 | 739422 | 1679 | 1247 | 1247 | 583 | 583 |
| | Damaliscus_lunatus | 6341937 | 6340372 | 5393454 | 15971 | 10942 | 10804 | 7947 | 7830 |
| | Hippotragus_niger | 4331751 | 4331187 | 4011095 | 12032 | 8116 | 8099 | 7611 | 7609 |
| | Redunca_redunca | 6171205 | 6170311 | 5025385 | 11074 | 7639 | 7627 | 7439 | 7403 |
| | Antidorcas_marsupialis | 15770605 | 15768692 | 11945184 | 24957 | 17765 | 17713 | 17301 | 17281 |
| | Litocranius_walleri | 11124101 | 11122087 | 8387053 | 23735 | 14910 | 14859 | 3063 | 2476 |
| | Eudorcas_thomsonii | 14262775 | 14251324 | 6489589 | 19774 | 13085 | 10102 | 7643 | 7486 |
| | Cephalophus_harveyi | 6957545 | 6954378 | 4913687 | 16155 | 11879 | 11767 | 9089 | 8962 |
| | Oreotragus_oreotragus | 13060574 | 13057358 | 9566685 | 19196 | 13865 | 13704 | 10804 | 10688 |
| | Bos_mutus | 4839035 | 4838378 | 4270063 | 12873 | 8515 | 8484 | 8254 | 8208 |
| | Syncerus_caffer | 20710227 | 20670757 | 18247129 | 52939 | 34999 | 26510 | 20026 | 19532 |
| | Tragelaphus_strepsiceros | 6216438 | 6215254 | 4996545 | 13346 | 9538 | 9516 | 8065 | 7965 |
| | Moschus_berezovskii | 15005110 | 14982232 | 9580723 | 25503 | 16825 | 11606 | 9768 | 9356 |
| | Capreolus_pygargus | 11261623 | 11260278 | 9510421 | 27361 | 18491 | 18478 | 18088 | 18079 |
| | Hydropotes_inermis | 6694661 | 6693334 | 5263569 | 16231 | 10985 | 10918 | 3346 | 3209 |
| | Odocoileus_virginianus | 13887384 | 13882602 | 12888814 | 32642 | 22651 | 22590 | 22147 | 21995 |
| | Rangifer_tarandus | 6752921 | 6751963 | 6235345 | 12112 | 8892 | 8892 | 8521 | 8520 |
| | Cervus_hanglu | 5210533 | 5209493 | 4747395 | 14482 | 9706 | 9685 | 9303 | 9298 |
| | Przewalskium_albirostris | 28139113 | 28132029 | 25944270 | 85893 | 59134 | 59092 | 56004 | 54210 |
| | Muntiacus_reevesi | 4111903 | 4111509 | 3631262 | 10967 | 7306 | 7301 | 3904 | 3848 |
| | Giraffa_tippelskirchi | 2390187 | 2389857 | 2106027 | 5510 | 4031 | 3928 | 2588 | 2584 |
| | Okapia_johnstoni | 3818826 | 3817344 | 3182923 | 9056 | 6724 | 6612 | 3840 | 3827 |
| | Tragulus_javanicus | 9572985 | 9572600 | 9157795 | 35322 | 24329 | 24329 | 23267 | 23253 |
| | Delphinapterus_leucas | 2084051 | 2083198 | 1866270 | 6740 | 4478 | 4455 | 4392 | 4387 |
| | Monodon_monoceros | 1229573 | 1229178 | 964606 | 3703 | 2472 | 2460 | 1985 | 1982 |
| | Neophocaena_asiaeorientalis | 3954634 | 3951830 | 3500255 | 11097 | 7397 | 4895 | 4655 | 4578 |
| | Phocoena_phocoena | 4242465 | 4241912 | 3806463 | 8581 | 6343 | 6343 | 6262 | 6260 |
| | Globicephala_melas | 1888740 | 1888453 | 1711254 | 5627 | 3849 | 3840 | 3475 | 3475 |
| | Sousa_chinensis | 1465607 | 1464423 | 1062513 | 3429 | 2225 | 1047 | 895 | 888 |

| | | | | | | | | | |
|---|---|---|---|---|---|---|---|---|---|
| 835 | Tursiops_truncatus | 3314718 | 3313772 | 3185866 | 9888 | 6538 | 3543 | 3494 | 3494 |
| | Lagenorhynchus_obliquidens | 2729131 | 2728839 | 2533884 | 8266 | 5569 | 5552 | 5197 | 5195 |
| | Orcinus_orca | 1502191 | 1501193 | 1171985 | 5265 | 2946 | 2822 | 2784 | 2780 |
| | Eschrichtius_robustus | 1683323 | 1682784 | 1311797 | 3226 | 2374 | 2374 | 2010 | 2010 |
| | Hippopotamus_amphibius | 6463386 | 6463116 | 4513924 | 8064 | 4879 | 4878 | 4762 | 4762 |
| | Camelus_ferus | 2785708 | 2784833 | 2337081 | 8135 | 5278 | 5277 | 5175 | 5155 |
| | Vicugna_pacos | 7676690 | 7671002 | 5841961 | 19031 | 12787 | 12017 | 11798 | 11796 |
| | Dicerorhinus_sumatrensis | 3559123 | 3558942 | 3192919 | 8901 | 6050 | 6049 | 5637 | 5587 |
| | Diceros_bicornis | 3852154 | 3851608 | 3087018 | 9619 | 6390 | 6335 | 655 | 627 |
| | Tapirus_terrestris | 7352432 | 7352142 | 7031044 | 17797 | 12657 | 12655 | 11798 | 11797 |
| | Manis_javanica | 6370097 | 6368844 | 5516276 | 14076 | 10012 | 9961 | 9548 | 9512 |
| | Ailuropoda_melanoleuca | 5814268 | 5809637 | 3766171 | 9177 | 5947 | 4229 | 4028 | 3966 |
| | Ursus_thibetanus | 6722212 | 6721541 | 6379954 | 20346 | 13840 | 13774 | 13520 | 13519 |
| | Ailurus_fulgens | 2935613 | 2934808 | 2550994 | 9292 | 6336 | 6317 | 6016 | 6003 |
| | Enhydra_lutris | 930695 | 930290 | 738019 | 2816 | 1829 | 1765 | 1657 | 1657 |
| | Lutra_lutra | 1789073 | 1788504 | 1521355 | 4930 | 3275 | 3195 | 2806 | 2741 |
| | Lontra_canadensis | 2305347 | 2305054 | 2102188 | 8110 | 5373 | 5344 | 4998 | 4998 |
| | Mustela_putorius | 6946880 | 6933923 | 6121941 | 34028 | 24489 | 2757 | 1274 | 1203 |
| | Neovison_vison | 4547129 | 4441163 | 3093604 | 8357 | 5291 | 4134 | 3508 | 3381 |
| | Martes_zibellina | 4975605 | 4973781 | 4447880 | 16159 | 10541 | 10472 | 10176 | 10149 |
| | Eumetopias_jubatus | 1118947 | 1118702 | 968190 | 3540 | 2331 | 2310 | 1978 | 1978 |
| | Odobenus_rosmarus | 2126365 | 2125178 | 1876071 | 7690 | 4682 | 4493 | 3957 | 3954 |
| | Halichoerus_grypus | 1983016 | 1982384 | 1222905 | 3119 | 2045 | 2001 | 1446 | 1437 |
| | Mirounga_leonina | 4172295 | 4172045 | 4054244 | 14006 | 9498 | 9493 | 8979 | 8972 |
| | Lycaon_pictus | 2106705 | 2105362 | 1688246 | 6549 | 4167 | 3779 | 2092 | 2053 |
| | Vulpes_vulpes | 4185824 | 4184669 | 3703756 | 9157 | 5926 | 5711 | 5547 | 5541 |
| | Crocuta_crocuta | 3391827 | 3388931 | 2881647 | 10556 | 7096 | 6506 | 6080 | 6059 |
| | Cryptoprocta_ferox | 2566789 | 2566564 | 2415647 | 6470 | 4712 | 4712 | 4323 | 4323 |
| | Helogale_parvula | 2175611 | 2175476 | 2059405 | 5131 | 3862 | 3860 | 2977 | 2977 |
| | Mungos_mungo | 4693201 | 4693082 | 4579866 | 13028 | 9406 | 9406 | 9117 | 9116 |
| | Suricata_suricatta | 11074252 | 11062917 | 10551371 | 38380 | 25701 | 16330 | 15615 | 15597 |
| | Paradoxurus_hermaphroditus | 7885209 | 7885086 | 7699389 | 14772 | 11188 | 11188 | 10174 | 10174 |
| | Acinonyx_jubatus | 1357617 | 1357341 | 1238946 | 4715 | 3155 | 3106 | 3025 | 3025 |
| | Panthera_tigris | 2490110 | 2486780 | 1934636 | 7975 | 5600 | 2826 | 2657 | 2656 |
| | Alouatta_palliata | 2109318 | 2108867 | 1590383 | 1547 | 1074 | 1073 | 737 | 732 |
| | Aotus_nancymaae | 12432061 | 12429888 | 11474271 | 28686 | 19078 | 19026 | 18684 | 18646 |
| | Cebus_imitator | 2683402 | 2680892 | 1992767 | 6817 | 4458 | 4317 | 4016 | 3999 |
| | Sapajus_apella | 4408932 | 4408079 | 4287174 | 11137 | 7493 | 7471 | 5727 | 5700 |
| | Saimiri_boliviensis | 2366752 | 2365351 | 2276134 | 8151 | 5347 | 5078 | 4491 | 4439 |
| | Callithrix_jacchus | 6982018 | 6976812 | 6436870 | 14788 | 9673 | 6706 | 6428 | 6405 |
| | Saguinus_imperator | 6940153 | 6939606 | 6915224 | 9486 | 7069 | 7069 | 6625 | 6623 |
| | Cercocebus_atys | 7231882 | 7227337 | 5885645 | 19716 | 11722 | 11207 | 4348 | 4280 |
| | Papio_anubis | 7305776 | 7301145 | 6468693 | 23092 | 15287 | 13444 | 3163 | 2880 |
| | Theropithecus_gelada | 2628228 | 2627470 | 2183835 | 5934 | 3992 | 3984 | 3822 | 3820 |
| | Mandrillus_leucophaeus | 6658637 | 6655822 | 5730681 | 13349 | 8931 | 8890 | 8557 | 8493 |
| | Macaca_nemestrina | 9845950 | 9837537 | 8694095 | 23707 | 15774 | 15614 | 15507 | 15490 |
| | Chlorocebus_sabeus | 6016217 | 5392043 | 4667211 | 11826 | 7670 | 7572 | 7455 | 7449 |
| | Colobus_angolensis | 4457083 | 4454753 | 3785790 | 11287 | 7356 | 7348 | 7103 | 7013 |
| | Pygathrix_nemaeus | 4252593 | 4252046 | 4190641 | 6750 | 4927 | 4927 | 4692 | 4686 |
| | Rhinopithecus_roxellana | 2326019 | 2325011 | 1875235 | 4565 | 2977 | 2042 | 1903 | 1893 |
| | Trachypithecus_francoisi | 4369518 | 4367252 | 4070721 | 11625 | 7482 | 5608 | 4975 | 4952 |
| | Gorilla_gorilla | 5681203 | 5679847 | 5040325 | 13577 | 8947 | 8924 | 7298 | 7285 |
| | Pan_troglodytes | 5139322 | 5135642 | 4589729 | 13282 | 8793 | 6694 | 4335 | 4323 |

| | | | | | | | | | |
|---|---|---|---|---|---|---|---|---|---|
| 836 | Pongo_abelii | 7447568 | 7445891 | 7047265 | 17979 | 11738 | 11661 | 10256 | 10236 |
| | Hylobates_moloch | 5911583 | 5911356 | 5262991 | 15500 | 10294 | 10294 | 8969 | 8945 |
| | Cheirogaleus_medius | 7652246 | 7652095 | 7455169 | 444243 | 333087 | 333087 | 78906 | 52463 |
| | Mirza_zaza | 3455259 | 3454926 | 3293758 | 7704 | 5513 | 5467 | 5285 | 5285 |
| | Lemur_catta | 10045555 | 10033595 | 9512017 | 25503 | 17584 | 11234 | 11055 | 11048 |
| | Prolemur_simus | 6459577 | 6457829 | 6066969 | 20307 | 13388 | 13376 | 13306 | 13305 |
| | Daubentonia_madagascariensis | 1470338 | 1470168 | 1361713 | 4811 | 3295 | 3295 | 3245 | 3241 |
| | Galeopterus_variegatus | 10155633 | 10155118 | 9439045 | 22523 | 14995 | 14995 | 13606 | 13602 |
| | Ochotona_princeps | 4125457 | 4123553 | 3518687 | 13886 | 9627 | 9597 | 9318 | 9286 |
| | Aplodontia_rufa | 5990566 | 5989663 | 5193284 | 7948 | 5990 | 5989 | 4515 | 4514 |
| | Cynomys_gunnisoni | 2962451 | 2961464 | 1743972 | 3601 | 2430 | 2411 | 2226 | 2189 |
| | Ictidomys_tridecemlineatus | 8372419 | 8363194 | 6245883 | 16012 | 10122 | 10081 | 7965 | 7933 |
| | Urocitellus_parryii | 5991789 | 5985440 | 3879617 | 11855 | 7766 | 7715 | 6668 | 6173 |
| | Spermophilus_dauricus | 6867570 | 6859530 | 4769752 | 13889 | 9337 | 6523 | 5954 | 5946 |
| | Marmota_himalayana | 3579162 | 2811360 | 1821584 | 4838 | 3245 | 3075 | 2797 | 2784 |
| | Xerus_inauris | 3216209 | 3215987 | 2990103 | 5764 | 4193 | 4193 | 2468 | 2468 |
| | Muscardinus_avellanarius | 523760 | 523679 | 386674 | 578 | 429 | 429 | 366 | 366 |
| | Acomys_cahirinus | 856064 | 855724 | 509494 | 2464 | 1528 | 1528 | 1160 | 1114 |
| | Meriones_unguiculatus | 1502162 | 1501971 | 1234032 | 2412 | 1745 | 1745 | 1489 | 1489 |
| | Psammomys_obesus | 3571352 | 3568492 | 2740708 | 11112 | 7517 | 7401 | 3927 | 3923 |
| | Mastomys_coucha | 2311275 | 2307558 | 988362 | 4242 | 2595 | 2590 | 1841 | 1708 |
| | Microtus_agrestis | 16092853 | 16063007 | 14704014 | 39199 | 25543 | 17721 | 13047 | 12976 |
| | Myodes_glareolus | 9773210 | 9548168 | 8387090 | 14023 | 8949 | 8782 | 8397 | 8242 |
| | Ondatra_zibethicus | 3597723 | 3597584 | 3415527 | 6925 | 5107 | 5107 | 4959 | 4958 |
| | Peromyscus_maniculatus | 6400141 | 6398315 | 5550936 | 18175 | 12776 | 12661 | 12282 | 12272 |
| | Sigmodon_hispidus | 699714 | 699415 | 372019 | 203 | 88 | 88 | 33 | 33 |
| | Cricetomys_gambianus | 3934123 | 3933512 | 3330123 | 8122 | 5995 | 5995 | 4660 | 4660 |
| | Nannospalax_galili | 7328757 | 7322576 | 5224805 | 9123 | 5957 | 5814 | 4862 | 4842 |
| | Rhizomys_pruinosus | 18346489 | 18317698 | 16326904 | 30593 | 20941 | 12915 | 4571 | 4347 |
| | Dipodomys_ordii | 2328055 | 2324478 | 1659079 | 4136 | 2799 | 2749 | 2702 | 2702 |
| | Cavia_tschudii | 2816202 | 2815610 | 2061480 | 3841 | 2781 | 2781 | 2212 | 2210 |
| | Hydrochoerus_hydrochaeris | 2084558 | 2084299 | 1770217 | 5668 | 3948 | 3948 | 2931 | 2928 |
| | Dasyprocta_punctata | 11616897 | 11616576 | 11341087 | 18602 | 13899 | 13899 | 12325 | 12324 |
| | Dinomys_branickii | 1329891 | 1329524 | 922130 | 2668 | 1980 | 1980 | 1394 | 1394 |
| | Fukomys_damarensis | 2172396 | 2171362 | 1682032 | 6303 | 4212 | 4146 | 3867 | 3862 |
| | Heterocephalus_glaber | 1828295 | 1827375 | 1309899 | 5287 | 3515 | 3388 | 3157 | 3087 |
| | Hystrix_cristata | 8063611 | 8063143 | 7527022 | 18754 | 13685 | 13684 | 10949 | 10945 |
| | Ctenodactylus_gundi | 3847485 | 3847306 | 3556424 | 10184 | 7033 | 7033 | 6841 | 6818 |

Whole_species_filtering_steps

| 837 | sp | accesion_nb | Busco+cov_filter | Ali+genus_filter | VCF_filter | Low_Pi_sp | 10Matree |
|---|---|---|---|---|---|---|---|
| | Acinonyx_jubatus | GCF_003709585.1 | pass | pass | pass | False | True |
| | Acomys_cahirinus | GCA_004027535.1 | pass | pass | removed | NA | True |
| | Aeorestes_cinereus | GCA_011751065.1 | pass | pass | pass | False | True |
| | Aepyceros_melampus | GCA_006408695.1 | pass | pass | pass | False | False |
| | Ailuropoda_melanoleuca | GCF_002007445.1 | pass | pass | pass | False | True |
| | Ailurus_fulgens | GCA_002007465.1 | pass | pass | pass | False | True |
| | Alouatta_palliata | GCA_004027835.1 | pass | pass | pass | True | True |
| | Ammotragus_lervia | GCA_002201775.1 | pass | pass | pass | False | False |
| | Anoura_caudifer | GCA_004027475.1 | pass | pass | pass | False | True |
| | Antidorcas_marsupialis | GCA_006408585.1 | pass | pass | pass | False | False |
| | Aotus_nancymaae | GCF_000952055.2 | pass | pass | pass | False | False |
| | Aplodontia_rufa | GCA_004027875.1 | pass | pass | pass | False | True |
| | Beatragus_hunteri | GCA_004027495.1 | pass | pass | pass | True | False |
| | Bos_mutus | GCA_007646595.3 | pass | pass | pass | False | True |
| | Callithrix_jacchus | GCA_009663435.2 | pass | pass | pass | False | True |
| | Camelus_ferus | GCF_009834535.1 | pass | pass | pass | False | True |
| | Capra_aegagrus | GCA_000765075.1 | pass | pass | pass | False | False |
| | Capreolus_pygargus | GCA_012922965.1 | pass | pass | pass | False | False |
| | Cavia_tschudii | GCA_004027695.1 | pass | pass | pass | False | True |
| | Cebus_imitator | GCF_001604975.1 | pass | pass | pass | False | True |
| | Cephalophus_harveyi | GCA_006410635.1 | pass | pass | removed | NA | False |
| | Cercocebus_atys | GCF_000955945.1 | pass | pass | pass | False | False |
| | Cervus_hanglu | GCA_010411085.1 | pass | pass | pass | False | False |
| | Cheirogaleus_medius | GCA_004024725.1 | pass | pass | removed | NA | True |
| | Chlorocebus_sabeus | GCF_000409795.2 | pass | pass | pass | False | False |
| | Choloepus_hoffmanni | GCA_000164785.2 | pass | pass | pass | False | True |
| | Colobus_angolensis | GCF_000951035.1 | pass | pass | pass | False | True |
| | Condylura_cristata | GCF_000260355.1 | pass | pass | pass | False | True |
| | Cricetomys_gambianus | GCA_004027575.1 | pass | pass | pass | False | True |
| | Crocuta_crocuta | GCA_008692635.1 | pass | pass | pass | False | True |
| | Cryptoprocta_ferox | GCA_004023885.1 | pass | pass | pass | False | True |
| | Ctenodactylus_gundi | GCA_004027205.1 | pass | pass | pass | False | True |
| | Cynomys_gunnisoni | GCA_011316645.1 | pass | pass | pass | False | False |
| | Damaliscus_lunatus | GCA_006408505.1 | pass | pass | pass | False | False |
| | Dasyprocta_punctata | GCA_004363535.1 | pass | pass | pass | False | True |
| | Daubentonia_madagascariensis | GCA_004027145.1 | pass | pass | pass | False | True |
| | Delphinapterus_leucas | GCF_002288925.2 | pass | pass | pass | False | True |
| | Desmodus_rotundus | GCF_002940915.1 | pass | pass | pass | False | True |
| | Dicerorhinus_sumatrensis | GCA_002844835.1 | pass | pass | pass | False | True |
| | Diceros_bicornis | GCA_004027315.2 | pass | pass | pass | True | True |
| | Dinomys_branickii | GCA_004027595.1 | pass | pass | pass | False | True |
| | Dipodomys_ordii | GCF_000151885.1 | pass | pass | pass | False | True |
| | Enhydra_lutris | GCF_002288905.1 | pass | pass | pass | False | False |
| | Eschrichtius_robustus | GCA_004363415.1 | pass | pass | pass | False | True |
| | Eudorcas_thomsonii | GCA_006408755.1 | pass | pass | pass | False | False |
| | Eumetopias_jubatus | GCF_004028035.1 | pass | pass | pass | False | True |
| | Fukomys_damarensis | GCF_012274545.1 | pass | pass | pass | False | True |
| | Galeopterus_variegatus | GCA_004027255.2 | pass | pass | pass | False | True |
| | Giraffa_tippelskirchi | GCA_001651235.1 | pass | pass | pass | False | True |
| | Globicephala_melas | GCF_006547405.1 | pass | pass | pass | False | False |
| | Gorilla_gorilla | GCF_008122165.1 | pass | pass | pass | False | False |
| | Halichoerus_grypus | GCA_012393455.1 | pass | pass | pass | False | True |

| | | | | | | | |
|---|---|---|---|---|---|---|---|
| 838 | Helogale_parvula | GCA_004023845.1 | pass | pass | pass | False | False |
| | Heterocephalus_glaber | GCF_000247695.1 | pass | pass | pass | False | True |
| | Hippopotamus_amphibius | GCA_004027065.2 | pass | pass | pass | False | True |
| | Hipposideros_armiger | GCF_001890085.1 | pass | pass | pass | False | True |
| | Hippotragus_niger | GCA_006942125.1 | pass | pass | pass | False | False |
| | Hydrochoerus_hydrochaeris | GCA_004027455.1 | pass | pass | pass | False | True |
| | Hydropotes_inermis | GCA_006459105.1 | pass | pass | pass | False | False |
| | Hylobates_moloch | GCF_009828535.2 | pass | pass | pass | False | True |
| | Hystrix_cristata | GCA_004026905.1 | pass | pass | pass | False | True |
| | Ictidomys_tridecemlineatus | GCF_000236235.1 | pass | pass | pass | False | False |
| | Lagenorhynchus_obliquidens | GCF_003676395.1 | pass | pass | pass | False | False |
| | Lemur_catta | GCA_004024665.1 | pass | pass | pass | False | True |
| | Litocranius_walleri | GCA_006410535.1 | pass | pass | removed | NA | False |
| | Lontra_canadensis | GCF_010015895.1 | pass | pass | pass | False | False |
| | Loxodonta_africana | GCF_000001905.1 | pass | pass | pass | False | True |
| | Lutra_lutra | GCA_902655055.1 | pass | pass | pass | False | True |
| | Lycaon_pictus | GCA_004216515.1 | pass | pass | pass | False | False |
| | Macaca_nemestrina | GCF_000956065.1 | pass | pass | pass | False | True |
| | Macroglossus_sobrinus | GCA_004027375.1 | pass | pass | pass | False | True |
| | Mandrillus_leucophaeus | GCF_000951045.1 | pass | pass | pass | False | False |
| | Manis_javanica | GCF_001685135.1 | pass | pass | pass | False | True |
| | Marmota_himalayana | GCA_005280165.1 | pass | pass | pass | False | True |
| | Martes_zibellina | GCA_012583365.1 | pass | pass | pass | False | True |
| | Mastomys_coucha | GCF_008632895.1 | pass | pass | removed | NA | True |
| | Megaderma_lyra | GCA_004026885.1 | pass | pass | pass | False | True |
| | Meriones_unguiculatus | GCA_004026785.1 | pass | pass | pass | False | True |
| | Micronycteris_hirsuta | GCA_004026765.1 | pass | pass | pass | False | True |
| | Microtus_agrestis | GCA_902806755.1 | pass | pass | pass | False | True |
| | Miniopterus_natalensis | GCF_001595765.1 | pass | pass | pass | False | True |
| | Mirounga_leonina | GCF_011800145.1 | pass | pass | pass | False | True |
| | Mirza_zaza | GCA_008750895.1 | pass | pass | pass | False | True |
| | Monodon_monoceros | GCF_005190385.1 | pass | pass | pass | False | False |
| | Moschus_berezovskii | GCA_006459085.1 | pass | pass | pass | False | True |
| | Mungos_mungo | GCA_004023785.1 | pass | pass | pass | False | False |
| | Muntiacus_reevesi | GCA_008787405.2 | pass | pass | pass | False | False |
| | Muscardinus_avellanarius | GCA_004027005.1 | pass | pass | pass | True | True |
| | Mustela_putorius | GCA_009859225.1 | pass | pass | pass | False | False |
| | Myodes_glareolus | GCA_004368595.1 | pass | pass | pass | False | False |
| | Nannospalax_galili | GCF_000622305.1 | pass | pass | pass | False | True |
| | Neophocaena_asiaeorientalis | GCF_003031525.1 | pass | pass | pass | False | True |
| | Neovison_vison | GCA_900108605.1 | pass | pass | pass | False | True |
| | Ochotona_princeps | GCF_000292845.1 | pass | pass | pass | False | True |
| | Odobenus_rosmarus | GCF_000321225.1 | pass | pass | pass | False | True |
| | Odocoileus_virginianus | GCF_002102435.1 | pass | pass | pass | False | True |
| | Okapia_johnstoni | GCA_001660835.1 | pass | pass | pass | False | False |
| | Ondatra_zibethicus | GCA_004026605.1 | pass | pass | pass | False | False |
| | Orcinus_orca | GCF_000331955.2 | pass | pass | pass | False | True |
| | Oreotragus_oreotragus | GCA_006410675.1 | pass | pass | pass | False | False |
| | Ovis_aries | GCA_000765115.1 | pass | pass | pass | False | True |
| | Pan_troglodytes | GCF_002880755.1 | pass | pass | pass | False | True |
| | Panthera_tigris | GCF_000464555.1 | pass | pass | pass | False | False |
| | Papio_anubis | GCF_008728515.1 | pass | pass | pass | False | False |
| | Paradoxurus_hermaphroditus | GCA_004024585.1 | pass | pass | pass | False | True |

| | | | | | | | |
|---|---|---|---|---|---|---|---|
| 839 | Peromyscus_maniculatus | GCF_000500345.1 | pass | pass | pass | False | True |
| | Phocoena_phocoena | GCA_004363495.1 | pass | pass | pass | False | False |
| | Phyllostomus_discolor | GCF_004126475.1 | pass | pass | pass | False | True |
| | Pongo_abelii | GCF_002880775.1 | pass | pass | pass | False | True |
| | Procavia_capensis | GCA_004026925.2 | pass | pass | pass | False | True |
| | Prolemur_simus | GCA_003258685.1 | pass | pass | pass | False | False |
| | Przewalskium_albirostris | GCA_006408465.1 | pass | pass | removed | NA | False |
| | Psammomys_obesus | GCA_002215935.2 | pass | pass | pass | False | False |
| | Pteropus_alecto | GCF_000325575.1 | pass | pass | pass | False | True |
| | Pygathrix_nemaeus | GCA_004024825.1 | pass | pass | pass | False | False |
| | Rangifer_tarandus | GCA_004026565.1 | pass | pass | pass | False | False |
| | Redunca_redunca | GCA_006410935.1 | pass | pass | pass | False | False |
| | Rhinolophus_ferrumequinum | GCF_004115265.1 | pass | pass | pass | False | True |
| | Rhinopithecus_roxellana | GCF_007565055.1 | pass | pass | pass | False | False |
| | Rhizomys_pruinosus | GCA_009823505.1 | pass | pass | pass | False | True |
| | Rousettus_aegyptiacus | GCF_001466805.2 | pass | pass | pass | False | True |
| | Saguinus_imperator | GCA_004024885.1 | pass | pass | pass | False | False |
| | Saimiri_boliviensis | GCF_000235385.1 | pass | pass | pass | False | True |
| | Sapajus_apella | GCF_009761245.1 | pass | pass | pass | False | False |
| | Sigmodon_hispidus | GCA_004025045.1 | pass | pass | pass | True | True |
| | Solenodon_paradoxus | GCA_004363575.1 | pass | pass | pass | False | True |
| | Sousa_chinensis | GCA_007760645.1 | pass | pass | pass | True | False |
| | Spermophilus_dauricus | GCA_002406435.1 | pass | pass | pass | False | False |
| | Suricata_suricatta | GCF_006229205.1 | pass | pass | pass | False | True |
| | Syncerus_caffer | GCA_902825105.1 | pass | pass | pass | False | False |
| | Tapirus_terrestris | GCA_004025025.1 | pass | pass | pass | False | True |
| | Theropithecus_gelada | GCF_003255815.1 | pass | pass | pass | False | False |
| | Tonatia_saurophila | GCA_004024845.1 | pass | pass | pass | False | True |
| | Trachypithecus_francoisi | GCF_009764315.1 | pass | pass | pass | False | False |
| | Tragelaphus_strepsiceros | GCA_006410795.1 | pass | pass | pass | False | False |
| | Tragulus_javanicus | GCA_004024965.2 | pass | pass | pass | False | True |
| | Trichechus_manatus | GCF_000243295.1 | pass | pass | pass | False | True |
| | Tursiops_truncatus | GCF_011762595.1 | pass | pass | pass | False | False |
| | Urocitellus_parryii | GCF_003426925.1 | pass | pass | pass | False | False |
| | Ursus_thibetanus | GCA_009660055.1 | pass | pass | pass | False | True |
| | Vicugna_pacos | GCA_000767525.1 | pass | pass | pass | False | True |
| | Vulpes_vulpes | GCF_003160815.1 | pass | pass | pass | False | True |
| | Xerus_inauris | GCA_004024805.1 | pass | pass | pass | False | True |
| | Castor_canadensis | GCF_001984765.1 | pass | removed | NA | NA | NA |
| | Connochaetes_taurinus | GCA_006408615.1 | pass | removed | NA | NA | NA |
| | Eubalaena_japonica | GCA_004363455.1 | pass | removed | NA | NA | NA |
| | Hemitragus_hylocrius | GCA_004026825.1 | pass | removed | NA | NA | NA |
| | Heterohyrax_brucei | GCA_004026845.1 | pass | removed | NA | NA | NA |
| | Macaca_fascicularis | GCA_012559485.1 | pass | removed | NA | NA | NA |
| | Macaca_mulatta | GCF_003339765.1 | pass | removed | NA | NA | NA |
| | Manis_pentadactyla | GCA_000738955.1 | pass | removed | NA | NA | NA |
| | Marmota_flaviventris | GCF_003676075.2 | pass | removed | NA | NA | NA |
| | Marmota_monax | GCA_901343595.1 | pass | removed | NA | NA | NA |
| | Mellivora_capensis | GCA_004024625.1 | pass | removed | NA | NA | NA |
| | Miniopterus_schreibersii | GCA_004026525.1 | pass | removed | NA | NA | NA |
| | Mirounga_angustirostris | GCA_004023865.1 | pass | removed | NA | NA | NA |
| | Mormoops_blainvillei | GCA_004026545.1 | pass | removed | NA | NA | NA |
| | Moschus_moschiferus | GCA_011751665.1 | pass | removed | NA | NA | NA |

| | | | | | | | |
|---|---|---|---|---|---|---|---|
| 840 | Muntiacus_muntjak | GCA_008782695.1 | pass | removed | NA | NA | NA |
| | Neomonachus_schauinslandi | GCF_002201575.1 | pass | removed | NA | NA | NA |
| | Neotragus_moschatus | GCA_006410615.1 | pass | removed | NA | NA | NA |
| | Noctilio_leporinus | GCA_004026585.1 | pass | removed | NA | NA | NA |
| | Odocoileus_hemionus | GCA_004115125.1 | pass | removed | NA | NA | NA |
| | Ovis_canadensis | GCA_004026945.1 | pass | removed | NA | NA | NA |
| | Pan_paniscus | GCF_013052645.1 | pass | removed | NA | NA | NA |
| | Panthera_leo | GCA_008795835.1 | pass | removed | NA | NA | NA |
| | Panthera_onca | GCA_004023805.1 | pass | removed | NA | NA | NA |
| | Peromyscus_californicus | GCA_007827085.2 | pass | removed | NA | NA | NA |
| | Peromyscus_eremicus | GCA_902702925.1 | pass | removed | NA | NA | NA |
| | Peromyscus_leucopus | GCF_004664715.1 | pass | removed | NA | NA | NA |
| | Physeter_catodon | GCF_002837175.2 | pass | removed | NA | NA | NA |
| | Pithecia_pithecia | GCA_004026645.1 | pass | removed | NA | NA | NA |
| | Pteronura_brasiliensis | GCA_004024605.1 | pass | removed | NA | NA | NA |
| | Pteropus_giganteus | GCA_902729225.1 | pass | removed | NA | NA | NA |
| | Rhinopithecus_bieti | GCF_001698545.1 | pass | removed | NA | NA | NA |
| | Scalopus_aquaticus | GCA_004024925.1 | pass | removed | NA | NA | NA |
| | Tapirus_indicus | GCA_004024905.1 | pass | removed | NA | NA | NA |
| | Tursiops_aduncus | GCA_003227395.1 | pass | removed | NA | NA | NA |
| | Ursus_arctos | GCF_003584765.1 | pass | removed | NA | NA | NA |
| | Ursus_maritimus | GCF_000687225.1 | pass | removed | NA | NA | NA |
| | Vulpes_lagopus | GCA_004023825.1 | pass | removed | NA | NA | NA |
| | Zalophus_californianus | GCA_009762305.1 | pass | removed | NA | NA | NA |
| | Artibeus_jamaicensis | GCA_004027435.1 | removed | NA | NA | NA | NA |
| | Ctenomys_sociabilis | GCA_004027165.1 | removed | NA | NA | NA | NA |
| | Dipodomys_stephensi | GCA_004024685.1 | removed | NA | NA | NA | NA |
| | Hipposideros_galeritus | GCA_004027415.1 | removed | NA | NA | NA | NA |
| | Lasiurus_borealis | GCA_004026805.1 | removed | NA | NA | NA | NA |
| | Macaca_fuscata | GCA_003118495.1 | removed | NA | NA | NA | NA |
| | Microcebus_tavaratra | GCA_008750935.1 | removed | NA | NA | NA | NA |
| | Microgale_talazaci | GCA_004026705.1 | removed | NA | NA | NA | NA |
| | Myrmecophaga_tridactyla | GCA_004026745.1 | removed | NA | NA | NA | NA |
| | Nasalis_larvatus | GCA_004027105.1 | removed | NA | NA | NA | NA |
| | Neotragus_pygmaeus | GCA_006410875.1 | removed | NA | NA | NA | NA |
| | Plecturocebus_donacophilus | GCA_004027715.1 | removed | NA | NA | NA | NA |
| | Taxidea_taxus | GCA_003697995.1 | removed | NA | NA | NA | NA |
| | Uropsilus_gracilis | GCA_004024945.1 | removed | NA | NA | NA | NA |

#### 5.12 Shrinkage model

### Gene-branch shrinkage model

Nicolas Lartillot


October 17, 2024

The gene-branch shrinkage model is fundamentally an approximate version of the model originally introduced in Gobbo et al (2020), using the mapping approximation (Romiguier et al, 2012). It is used in the present context to detect outlier genes, i.e. genes that deviate in their synonymous branch lengths, compared to the average over all genes, and are thus suspected to be data errors (due to alignment errors or incorrect orthology assignment).

This model expresses the distribution of synonymous branch lengths and $dN/dS$ across genes and branches as a combination of a branch effect, a gene effect, and a residual effect, following a Gamma-Poisson structure. Thus, for gene $i$ on branch $j$, the local (gene-specific) branch length $l_{ij}$ and $dN/dS$ $\omega_{ij}$ are assumed to have a gamma prior:

$$
\begin{aligned}
l_{ij} &\sim Gamma(a_i b_j, \alpha) \\
\omega_{ij} &\sim Gamma(u_i v_j, \beta)
\end{aligned}
$$

where:

- $a_i$ and $b_j$ are the gene and branch effects on $dS$
- $u_i$ and $v_j$ are the gene and branch effects on $dN/dS$
- $\alpha$ and $\beta$ tune the variance of the residual gene-branch deviations

In a hierarchical Bayes framework, the priors on $a$, $b$, $u$, $v$ are all Gamma, with hyperpameters (mean and shape parameter) being themselves estimated. To make the model identifiable, the prior has mean of 1 for the $a_i$'s and for the $v_j$'s. With this convention, the mean over the $b_j$'s represents the mean synonymous length of branch $j$, while the mean over the $u_i$'s represents the mean $dN/dS$ for gene $i$.

This hierarchical model can then be combined with the Poisson likelihood for the mapping statistics:

$$K_{ij}^S \sim Poisson(l_{ij}\, L_{ij}^S) \tag{1}$$

$$K_{ij}^N \sim Poisson(l_{ij}\, \omega_{ij}\, L_{ij}^N) \tag{2}$$

Here, $K_{ij}^S$ and $K_{ij}^N$ are the synonymous and non-synonymous counts, for gene $i$ and branch $j$, and $L_{ij}^S$ and $L_{ij}^N$ are the corresponding numbers of mutational targets. These Poisson likelihoods can be seen as a link function, such that the overall procedure can be seen as a Bayesian hierarchical Poisson regression model, which, based on the 'observed' counts $K$ and covariates $L$, estimates gene- and branch-effects on $dS$ and $dN/dS$. It can also be seen as a Bayesian shrinkage device, in the sense that the gene-branch parameters $l_{ij}$ and $\omega_{ij}$ are shrunk towards their expected value (resp. $a_i b_j$ and $u_i v_j$) based on information-sharing in the two dimensions.

Shrinkage is particularly useful in the present context, as it will smooth out the stochastic errors on those gene-branch configurations that are characterized by low synonymous and non-synonymous counts (small gene length and/or short branch). As a result, and conversely, gene-branch configurations that in the end do show a large deviation between their posterior estimate for $l_{ij}$ and the expectation based on other genes and branches, $a_i b_j$, can only do so by virtue of a strong empirical signal. Thus, the shrinkage model efficiently filters out stochastic errors, so as to more clearly single out the statistically significant outliers.

This model was implemented in a simple MCMC framework. After running the MCMC, for a given gene $i$ and branch $j$, the $dS$ and $dN/dS$ deviations, defined as:

$$\begin{aligned} x_{ij} &= \frac{l_{ij}}{a_i b_j} \\ y_{ij} &= \frac{\omega_{ij}}{u_i v_j} \end{aligned}$$

were averaged over the MCMC sample, giving posterior mean estimates of the deviations. Genes with a $dS$ deviation larger than 3 were discarded.
